# Recording true oxygen reduction capacity during photosynthetic electron transfer in Arabidopsis thylakoids and intact leaves

**DOI:** 10.1101/2021.10.28.466325

**Authors:** Duncan Fitzpatrick, Eva-Mari Aro, Arjun Tiwari

**Author notes:** The author responsible for contact and ensuring the distribution of materials integral to the findings presented in this article in accordance with the Journal policy described in the Instructions for Authors (http://www.plantphysiol.org) are: Eva-Mari Aro,; Arjun Tiwari. **Footnotes**. Authors Contribution: AT, DF, E-MA designed research; AT, DF performed research; AT, DF, E-MA analysed data; DF, AT, E-MA, wrote the paper. **Competing interests:** Authors declare no competing interest.

## Abstract

Reactive oxygen species (ROS) are generated in electron transport processes of living organisms in oxygenic environments. Chloroplasts are plant bioenergetics hubs where imbalances between photosynthetic inputs and outputs drive ROS generation upon changing environmental conditions. Plants have harnessed various site-specific thylakoid membrane ROS products into environmental sensory signals. Our current understanding of ROS production in thylakoids suggests that oxygen (O_2_) reduction takes place at numerous components of the photosynthetic electron transfer chain (PETC). To refine models of site-specific O_2_ reduction capacity of various PETC components in isolated thylakoids, the stoichiometry of oxygen production and consumption reactions, associated with H_2_O_2_ accumulation, was quantified using membrane inlet mass spectrometry and specific inhibitors. Combined with P700 spectroscopy and electron paramagnetic resonance spin trapping, we demonstrate that electron flow to PSI is essential for H_2_O_2_ accumulation during light-induced photosynthetic electron transport process. Further leaf disc measurements provided clues that H_2_O_2_ from PETC has a potential of increasing mitochondrial respiration and CO_2_ release.

**One sentence summary:** Photosynthetically derived H_2_O_2_ only accumulates at Photosystem I and may trigger cooperation with mitochondria during stress

## Introduction

Plants exploit a range of redox balancing and signaling mechanisms to maintain an energetic homeostasis in chloroplasts, particularly important under fluctuating light conditions. These mechanisms alleviate over-excitation and over-reduction of critical photosynthetic electron transfer chain (PETC) components that may generate reactive oxygen species (ROS). Although ROS formation within the photosynthetic apparatus contributes to photodamage, they also function as critical feedback mechanisms and chemical messengers to modify and restore an optimal redox balance. Because ROS formation occurs at the crossroads between photosynthetic damage and regulation, the elucidation of the specific locations and mechanisms of photosynthetically derived ROS generation is critical for understanding the interactions between photodamage and photoprotection. Such knowledge is central for future projects aimed at improving photosynthetic productivity.

Understanding site specific ROS formation within the thylakoid membrane is complicated by the fact that the various components of the PETC participate in several redox reactions, forming a variety of site-specific ROS molecules with varying reactivity and reaction products. For example, during conditions of over-excitation, highly reactive and long-lived chlorophyll triplet states can form and photosensitize molecular oxygen (O_2_) to singlet oxygen (^1^O_2_) (Halliwell, Gutteridge 1984). This short-lived and highly reactive form of oxygen is mostly associated with PSII (Durrant, Giorgi et al. 1990, Telfer, Oldham et al. 1999) and leads to rapid peroxidation of proteins, lipids and nucleotides. On the other hand, electron transfer within the PETC can result in the reduction of O_2_ to form the superoxide anion (O_2_^−•^). This reaction was originally described by Mehler (Mehler 1951) and occurs primarily at the acceptor side of PSI (Mehler 1951, Furbank, Badger 1983). To counter this reaction, the catalyst superoxide dismutase (SOD) enzymatically dismutates two O_2_^−•^ molecules into the oxidant H_2_O_2_, with a release of one O_2_. Although H_2_O_2_ is relatively stable in biological systems, further reduction of H_2_O_2_ can result in the formation of highly destructive hydroxyl radicals (HO^•^). To minimise potential damage from such destructive molecules, strong ROS scavenging systems comprising peroxiredoxins, ascorbate and ascorbate peroxidase enzymes (Asada 2006) operate in both the thylakoid membrane and in the stroma. They facilitate the complete enzymatic two-electron reduction and protonation of H_2_O_2_, thereby yielding two water molecules and completing an electron pathway described as the water-water cycle (WWC) (Asada 1999). Despite the antioxidant systems that can efficiently quench ROS, a growing body of evidence suggests that photosynthetically derived ROS such as H_2_O_2_ act in environmental sensing (Mubarakshina, Ivanov 2010) and intercellular relay of information (Fichman, Miller et al. 2019) or function directly as a retrograde signal exported from the chloroplasts to the nucleus (Gollan, Aro 2020, Exposito-Rodriguez, Laissue et al. 2017).

Numerous strategies to measure site-specific ROS formation in PETC have been developed and applied over several decades. The resulting literature suggests that in addition to the Mehler reaction at PSI, O_2_^−•^ and H_2_O_2_ can form within the PETC at heavily reduced PSII (Tiwari, Pospisil 2009), the PQ pool (Khorobrykh, S. A., Ivanov 2002, Mubarakshina, Ivanov 2010, Mubarakshina Borisova, Kozuleva et al. 2012, Khorobrykh, Sergey A., Karonen et al. 2015), plastid terminal oxidase (PTOX) (Heyno, Gross et al. 2009) and Cyt-*b*_*6*_*f* complex (Baniulis, Hasan et al. 2013). The current work aimed at understanding whether these different sites can form stable H_2_O_2_ pools capable of functioning as secondary messengers in guiding plant acclimation according to the environmental cues. To this end, we measured the true capacity of oxygen reduction/consumption at PSII, PQ-pool, and Cyt-*b6f* in proportion to the PSII water oxidation rates.

### A new approach to site-specific ROS measurements

It is necessary to revisit site-specific ROS formation in the thylakoid membrane, incorporating stronger controls, higher precision and with the capacity to discriminate O_2_ reduction and true H_2_O_2_ accumulation. Incorporating lessons from our recent finding of the failure of DNP-INT to completely block electron transfer to PSI (Fitzpatrick, Aro et al. 2020), we have taken a methodical three-step approach to re-examine ROS formation and accumulation in isolated thylakoids of *Arabidopsis thaliana* (hereafter Arabidopsis) and have then tested our conclusions in-vivo.

As the first step, to avoid artefacts from any residual activity of the PSI-Mehler reaction, we measured P700 redox kinetics under experimental conditions to verify that each PETC inhibitor (Fig. 1, as described schematically) completely blocked the re-reduction of oxidized P700 (P700^+^). As the second step, we took advantage of the fact that isolated thylakoids lack the stroma and terminal electron acceptor NADP^+^, and thus the electrons derived from oxidation of H_2_O at PSII are transferred terminally to O_2_ via the PETC. The overall process can be dissected into partial reactions (Asada 2006) (Fig. 1, Reactions 1 – 4). These partial reactions can be estimated from the stoichiometry of O_2_ produced and O_2_ consumed at each step. Reaction (1) -The oxidation of two H_2_O molecules at PSII releases one O_2_ and four electrons into the PETC. Reaction (2) –The four electrons reduce four O_2_ molecules to O_2_^−•^. Reaction (3) - The four O_2_^−•^ dismutate or decompose (rapidly) into two H_2_O_2_ molecules, reforming two molecules of O_2_. Reaction (4) - A complete degradation of two H_2_O_2_ further yields one molecule of O_2_ and two H_2_O, the latter reforming the H_2_O originally oxidized at PSII, resulting in a net O_2_ flux of zero. At each of these 4 steps the stoichiometric ratio of O_2_ produced to O_2_ consumed varies, which we describe as the O_2_ flux ratio (Fig. 1, O_2_ Flux Ratios). The O_2_ flux ratio was quantified with MIMS by separating the two events from each other *i*.*e*. the rate of PSII H_2_O splitting reactions were measured by the production rate of the naturally abundant O_2_ isotopologue (Fig. 1, ^16^O_2_, blue font), whilst O_2_ reduction reactions were determined by the consumption rate of the artificially enriched heavy O_2_ isotopologue (Fig. 1, ^18^O_2_, red font).

**Figure 1.**
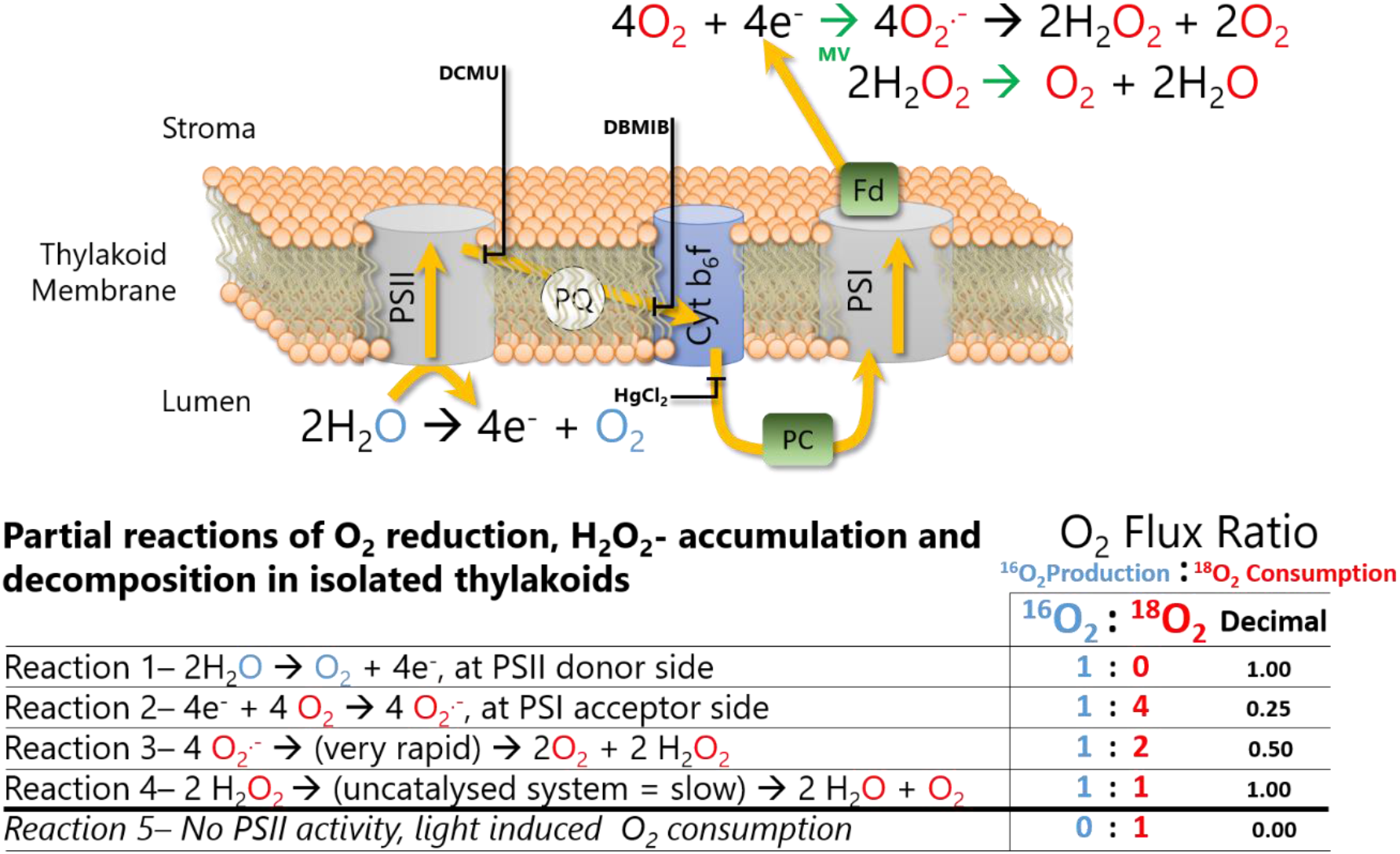
Schematic description of O_2_ reduction, H_2_O_2_ accumulation and H_2_O_2_ decomposition cycle operating in isolated thylakoid membrane samples, highlighting labelled isotope reactions and the targets of specific PETC inhibitors, and catalysts, used in this study. Water oxidation at PSII generates a ^16^O_2_, represented in blue font, and releases four electrons into the PETC, represented by yellow arrows. These four electrons reduce four artificially enriched ^18^O_2_, represented in red font. According to the reactions presented at the top right, the superoxide is dismutated into H_2_O_2_ and finally back to H_2_O, reproducing the H_2_O originally oxidized but now containing a red, labelled O_2_. This final reaction can be catalysed by externally added cat. (Catalase) in the isolated thylakoids, which instantly pushes the cycle to the end. Breaking the complete cycle into partial reactions (Reactions 1 to 4) provides a stoichiometric gas flux ratio to correlate each step of the cycle back to the O_2_ production to consumption ratio being measured with MIMS. Reactions 1 and 4 have the same ratio but are distinguished by the O_2_ uptake component, occurring only with artificial acceptors. Reaction 5 is a variant of the ratio, predicted from observations that DCMU blocks all water oxidation but samples were still able to consume O_2_ in a light dependent manner, suggesting ^1^O_2_ or organic peroxide formation.

The key advantages of the first and second steps of our approach, compared to measuring net O_2_ fluxes polarographically, are that (1) no assumptions are required to determine the number of electrons entering the PETC, (2) the 1:1 ratio of Reaction 4 is measurable, even if H_2_O_2_ is catalytically decomposed and (3) it is possible to clearly observe when O_2_ was consumed in the absence of electron transport, *i*.*e*. to observe the 0:1 stoichiometry described in Reaction 5 (Fig. 1), which results from lipid/protein peroxidation reactions such as those resulting from ^1^O_2_ formation. As the third step in our approach, Electron Paramagnetic Resonance (EPR) spin-trapping was applied to thylakoid samples under similar experimental conditions. This enabled us to directly infer the ROS formed at each location in the PETC, to support the observations and conclusions made with the MIMS data.

## Results

### Controlling for ROS artefacts resulting from only partial inhibition of PSI

As described in Figure 1, application of specific inhibitors enabled systematic segmentation of the PETC into discrete components. It was then possible to measure the site-specific ROS formation of each component with MIMS or EPR. However, to avoid artefacts from an active PSI-Mehler reaction during sample illumination, the efficacy of each inhibitor to completely block re-reduction of oxidised P700 (P700^+^, special chlorophyll pair in PSI) was measured via P700 spectroscopy with a Dual PAM, under emulated experimental conditions. The redox measurement and quantification of P700 in Dual PAM is based on strong absorption characteristics of oxidised P700 between 800 to 840 nm. A difference of transmittance between the 875 nm and 830 nm in darkness shows completely reduced P700. The calibrated kinetic signal of P700 is represented as ΔI/I×10^−3^ units. The following inhibitors were tested and used: (1) DCMU, as an inhibitor of PQ pool reduction, enabled PSII to be heavily reduced (Witt, Rumberg et al. 1968), (2) DBMIB, as an inhibitor of PQ oxidation by *Cyt-b*_*6*_*f*, enabled complete reduction of PSII and the PQ pool (Bauer, Wijnands 1974) and (3) HgCl_2_, as an inhibitor of plastocyanin (PC) function, enabled reduction of PSII, PQ pool and Cyt*-b*_*6*_*f* (Kimimura, Katoh 1972). Historically, DNP-INT has been used to block reduction of Cyt-*f* whilst enabling interaction of the low potential chain of Cyt-*b*_*6*_*f* complex with plastosemiquinone, considered necessary for the formation of O_2_^−•^ within the PQ-pool (Khorobrykh, S. A., Ivanov 2002, Mubarakshina, Ivanov 2010). However, in an attempt to maintain such dynamics between the PQ pool and Cyt*-b*_*6*_*f* in isolated thylakoids and leaf discs, we observed that DNP-INT failed to completely block the reduction of PSI (Fitzpatrick, Aro et al. 2020). Hence, HgCl_2_ was considered the best available alternative. Although HgCl_2_ has been reported to have multiple inhibitory effects on photosynthetic apparatus (Carpentier 2001), we observed strong PSII activity in isolated thylakoid membranes in the presence of HgCl_2_ under both growth light and high light (see Supplemental Fig. S1). Indeed, to avoid measuring artefacts with EPR or MIMS from a still active PSI, we systematically confirmed that each inhibitor could completely block reduction of oxidised PSI (achieved through application of a background FR light) from receiving electrons sourced at PSII following the application of strong actinic light pulses (ST and MT) (Fig. 2).

**Figure 2.**
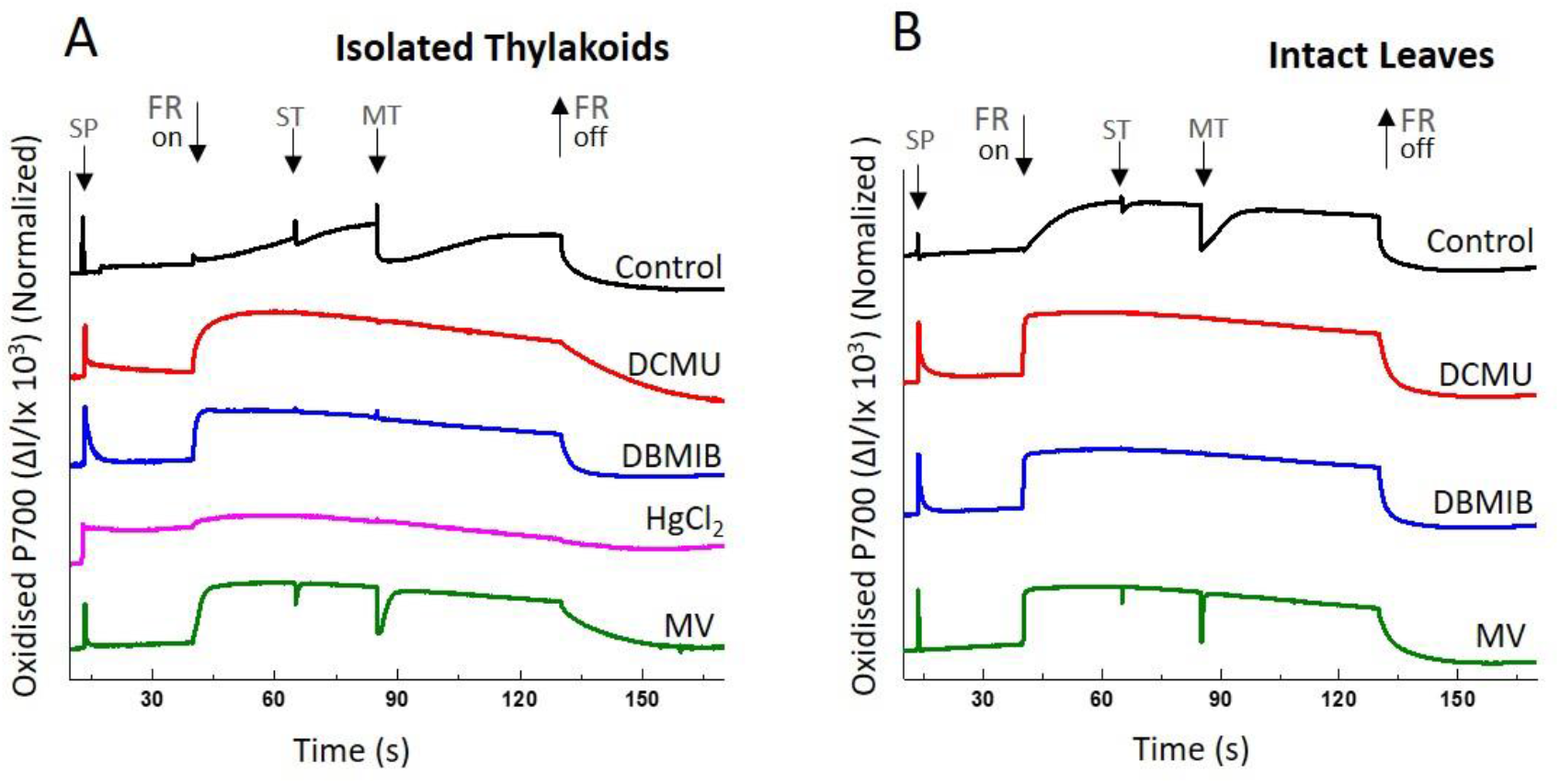
Comparison of site-specific inhibitors in blocking electron flow from PSII to PSI. Dark-adapted isolated thylakoids, or intact leaves, were exposed to a saturating pulse (SP) of actinic light. Following this, P700 re-reduction was measured under far red (FR) background illumination by firing strong actinic light pulses of 50 μs (single turnover, ST) and 50 ms (multiple turnover, MT) on A, isolated thylakoids equivalent to chlorophyll concentration of 80 μg chl ml^-1^ in measuring buffer containing the uncoupler NH_4_Cl (5mM) and B, intact leaves. Leaves were infiltrated in darkness using water for the control sample. Inhibitors/modulators were used at 10 μM concentration, except for HgCl_2_ used at 2 mg ml^-1^. Representative curves averaged from minimum 3 to 5 biological replicates.

Freshly prepared isolated thylakoid (Fig. 2A) and intact leaf (Fig. 2B) samples from Arabidopsis were incubated in darkness with 10 μM DCMU, 10 μM DBMIB or 2 mg ml^-1^ HgCl_2_ (we found HgCl_2_ toxic to intact leaves so this inhibitor was applied only for isolated thylakoid samples). As negative controls for blocking PSI activity, 10 μM Methyl Viologen (MV, catalyst of O_2_ reduction at PSI acceptor side) and untreated control samples were also measured. During the preceding dark incubation, a single saturating actinic light pulse (SP) revealed an overall capacity of each sample to rapidly oxidise P700 followed by fast re-reduction in all samples except for the HgCl_2_-treated thylakoids. Subsequent continuous background FR light (128 μmol photons m^-2^ s^-1^) preferentially oxidized P700 to P700^+^. To activate electron flow from PSII to PSI, a 50 μs single turnover (ST), followed by a 50 ms multiple turnover (MT) pulse of actinic light (10 000 μmol photons m^-2^ s^-1^) was superimposed over the FR background. The electrons released from water by PSII into the PETC by these actinic light pulses transiently reduced P700^+^ in both sets of untreated control samples (Fig. 2, A and B, black traces), evidenced by the pulse length dependent decrease, or ‘dip’ of the P700^+^ signal, before being fully re-oxidised by the continuous FR background light (Tiwari, Mamedov et al. 2016). In contrast, no transient dip was observed in the P700^+^ signal when thylakoid samples were treated with DCMU, DBMIB or HgCl_2_ (Fig. 2A, red, blue and pink color traces), or in leaves treated with DCMU or DBMIB (Fig. 2B, red and blue traces). This confirmed that these inhibitors blocked all electron transport from PSII to PSI. Uniquely, HgCl_2_ blocked the re-reduction of P700^+^ in isolated thylakoid samples following the actinic light pulse in the initial dark phase of the measurement (Fig. 2A, pink trace). This was consistent with HgCl_2_ targeting the function of PC, which was then severely restricted in its capacity to deliver electrons to P700^+^ throughout the rest of the measurement. Samples infiltrated with MV (Fig. 2, A and B, green traces) exhibited clear actinic light dependent dips. However, due to the accelerated rate of re-oxidation produced by the artificial PSI acceptor, and catalyst of O_2_^−•^ formation, the ‘dips’ following ST and MT pulses were ‘narrowed’ compared to the control samples (Fig. 2, A and B, black traces,) in which electrons took longer to find an acceptor.

### O_2_ flux ratio measured with MIMS positively identified H_2_O_2_ accumulation in thylakoid samples

Freshly prepared thylakoid samples were illuminated at a relatively low light intensity, equivalent to growth light (GL) (120 μmol photons m^-2^ s^-1^), to minimise ^1^O_2_ formation and to verify that H_2_O_2_ accumulation results in a 1:2 O_2_ flux ratio (Fig. 1) in MIMS measurements (Fig. 3, A-G). To ensure that the O_2_ flux ratios were calculated based on light dependent O_2_ fluxes, the background dark rate of O_2_ consumption (also observed in (Furbank, Badger 1983)) was offset to zero. Control samples (Fig. 3A) produced O_2_ in a relatively stable manner across the three minutes of illumination, at a rate of approximately 2.5 μmol mg chl^-1^ h^-1^. The simultaneous rate of O_2_ consumption was less stable, exhibiting a steady decrease in the integrated rate over the three minutes of illumination. However, the initial O_2_ consumption rate of approximately 5.0 μmol mg chl^-1^ hr^-1^ was double the initial O_2_ production rate, resulting in an O_2_ flux ratio of 1:2, consistent with accumulation of H_2_O_2_ (Fig. 1). The addition of catalase (700 unit per ml^-1^) to rapidly degrade any H_2_O_2_ formed did not drastically affect the rate or stability of O_2_ production (Fig. 3B). However, it decreased the absolute rate of O_2_ consumption so that it matched the O_2_ evolution rate, resulting in the 1:1 O_2_ flux ratio expected in the absence of H_2_O_2_ accumulation (Fig. 1). It is clear that when H_2_O_2_ accumulates, the rate of O_2_ consumption is greater than the rate of O_2_ evolution (Fig. 3, A and C) (red line is a greater magnitude than the blue line and net rate shown with a grey line is negative). In contrast, when the accumulation of H_2_O_2_ is excluded by the addition of catalase (Fig. 3, B and D) or in samples excluding PSI function (Fig. 3, E-G), we clearly observed that O_2_ consumption cannot significantly exceed O_2_ production, as any H_2_O_2_ formed was immediately decomposed, re-releasing the ^18^O_2_.

**Figure 3.**
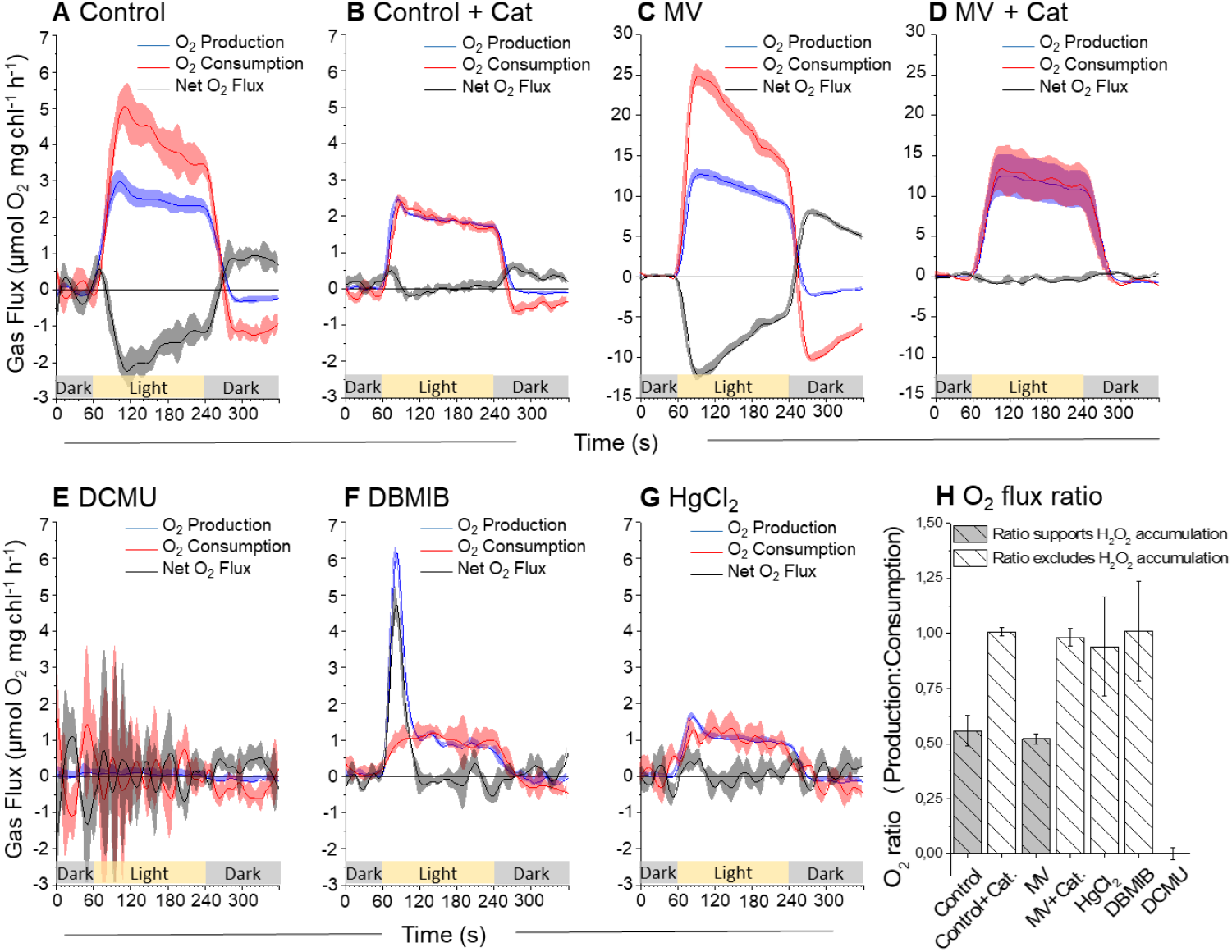
Integrated rates of ^16^O_2_ production and ^18^O_2_ consumption by isolated thylakoid samples, measured simultaneously at 120 μmol photons m^-2^ s^-1^ with MIMS. Illumination of samples represented by yellow bar with grey representing darkness. Curves A, untreated control, B, untreated control + catalase, C, 10 μM MV, D, 10 μM MV + catalase (observe increased scale for c and d), E, 10 μM DCMU, F, 10 μM DBMIB and G, 2 mg ml^-1^ HgCl_2_ are the product of averaging minimum three replicates plotted with standard error. H, The ratios of O_2_ production to consumption rates were calculated as described in text and plotted as a decimal value to highlight conditions in which H_2_O_2_ could accumulate, based on the stoichiometry of partial reactions from O_2_ reduction to water formation again. All measurements were performed in measurement buffer containing the uncoupler NH_4_Cl (5mM).

As a positive control for H_2_O_2_ accumulation, and to simultaneously test the sample’s acceptor side limitation at PSI, 10 μM MV was added to thylakoid samples. The O_2_ evolution rate increased to 12.5 μmol O_2_ mg chl^-1^ h^-1^ and the initial O_2_ consumption rate of 25.0 μmol O_2_ mg chl^-1^ h^-1^ reproduced the O_2_ flux ratio of 1:2 observed in control samples (Fig. 3C, observe change in y-axis scale). This result verified that control samples illuminated at GL were PSI acceptor limited, therefore all components of the thylakoid membrane were heavily reduced during illumination. The addition of catalase to MV-treated thylakoid samples (Fig. 3D) replicated the effects of catalase added to control thylakoids, reproducing a 1:1 O_2_ flux ratio with no effect on the rate of O_2_ production. The strong negative O_2_ consumption rate following illumination, both in control and amplified in MV treated samples, suggested that the declining O_2_ consumption rate during illumination was a product of ^18^O_2_ release from an accumulated pool of H_2_O_2_. This was supported by the observation that the sum of the O_2_ consumption rate at the end of the light period and the negative rate at the beginning of the dark period approximated the initial O_2_ consumption rate in untreated control and MV samples (for a more detailed explanation of interpreting time resolved gas flux transients, and how the negative O_2_ consumption burst has been interpreted as the decomposition of an accumulated H_2_O_2_ pool, refer to Supplemental Fig. S2). In the summary bar chart of the O_2_ flux ratios (Fig. 3H) it can be seen that the control and MV treated samples exhibited an O_2_ flux ratio of 1:2. Conversely, in samples where catalase precluded H_2_O_2_ accumulation an O_2_ flux ratio of 1:1 was observed. Hence, MIMS was able to discriminate between isolated thylakoid samples in which H_2_O_2_ was accumulated from those in which the H_2_O_2_ was rapidly decomposed.

### MIMS confirms that O_2_ reduction at PQ-pool and Cyt-*b*_*6*_*f* cannot accumulate H_2_O_2_

To compare the site-specific capacity of heavily reduced PETC components upstream of PSI to generate ROS and accumulate a pool of H_2_O_2_ in-vitro, we measured the O_2_ flux ratios of freshly isolated thylakoid samples incubated with 10 μM DCMU, 10 μM DBMIB or 2 mg ml^-1^ HgCl_2_. Based on the results of MV addition to isolated thylakoids, discussed above, we were confident that components within the PETC were heavily reduced at GL irradiance. Incubation with DCMU clearly impaired all O_2_ production (Fig. 3E) and although noisy, O_2_ consumption was minimal. In contrast, samples incubated with DBMIB exhibited strong O_2_ production by water oxidation at the initiation of illumination, in the absence of commensurate O_2_ consumption. This resulted in an initial O_2_ flux ratio of 1:0 (Fig. 1, Reaction 1), consistent with DBMIB accepting electrons from PSII as reported previously (Lozier, Butler 1972). The rate of O_2_ production peaked, presumably as available DBMIB was fully reduced, and then slowed as O_2_ consumption became the primary electron sink (more evidence that samples were acceptor-limited). The steady-state rate of O_2_ production and consumption of approximately 1.0 μmol O_2_ mg chl^-1^ h^-1^ (Fig. 3F) comprised an O_2_ flux ratio of 1:1. This ratio matched the addition of catalase to control and MV treated thylakoid samples. According to the results of a DBMIB concentration response curve, the integrated area of the O_2_ evolution peak was DBMIB concentration dependent (r^2^ of 0.99) whilst the steady-state rate following the peak was DBMIB concentration independent (Supplemental Fig. S3). This result thus suggested that DBMIB was not acting as a redox intermediate in the steady-state O_2_ consumption pathway. Incubating thylakoids with HgCl_2_ (Fig. 3G) produced a very similar PSII activity as that in DBMIB samples, exhibiting a steady-state rate of O_2_ production approximately 1.0 μmol O_2_ mg chl^-1^ h^-1^ and a 1:1 O_2_ flux ratio. Significantly, the lack of a large initial peak in O_2_ production supported the conclusions drawn from the DBMIB concentration response curve (Supplemental Fig. S3). The small initial peak in O_2_ production observed with HgCl_2_ was also evident in control samples with and without catalase (Fig. 3, A and B) but absent from DCMU curves, suggesting it was the product of reducing a small intrinsic pool of acceptors between PSII and PSI, likely the PQ pool.

The fact that HgCl_2_ and DBMIB samples exhibited almost identical steady-state rates and a 1:1 O_2_ flux ratio at GL supported the contention that a pathway to O_2_ reduction operates within the thylakoid membrane. Whilst this pathway upstream of PSI exhibited a smaller capacity to reduce O_2_ compared to the PSI-Mehler reaction, it clearly supports PSI independent steady-state turnover of PSII. PTOX may be a candidate to explain such a pathway, reducing O_2_ to H_2_O with reductant from PQH_2_ (Cournac, Josse et al. 2000). In an effort to control for PTOX function we repeated DBMIB measurements in the presence of the widely used PTOX inhibitors *n*-propyl gallate or octyl gallate (Cournac, Josse et al. 2000). Unexpectedly in our measurements, both treatments enhanced O_2_ production (PSII activity) and O_2_ consumption, even in the presence of DCMU (Supplemental Fig. S4, with short discussion of these experiments). We did not investigate this result further, concluding that based on analysis of the P700 kinetics of isolated thylakoids (Fig. 2A) and the O_2_ Flux ratios of thylakoid samples (Fig. 3H), H_2_O_2_ can only accumulate in the thylakoid membrane if electrons are able to reach PSI.

### Increasing the redox and excitation pressure highlighted separate ROS pathways within the thylakoid membrane in-vitro

To test the impact of increased excitation and redox pressure on the results described above, we repeated all MIMS measurements with inhibitors using high light illumination (HL, 900 μmol photons m^-2^ s^-1^ (Fig. 4). The untreated control samples increased rates of O_2_ production and consumption to approximately 5 and 10 μmol O_2_ mg chl^-1^ h^-1^,respectively (Fig. 4A), whilst the 1:2 O_2_ flux ratio was maintained. We assured that relatively low rates observed were a result of the lack of electron acceptors in isolated thylakoids (no externally added ferredoxin (Fd)), and the absence of extra SOD in our experiments. Contrasting to untreated control thylakoids, the DCMU response was markedly different at higher illumination. The dosage required to fully impair PSII function increased to 50 μM and a strong light dependent, catalase independent, O_2_ consumption was observed at a rate of approximately 1.5 μmol O_2_ mg chl^-1^ h^-1^ (Fig. 4, B and F), consistent with previously published observations (Khorobrykh, S. A., Khorobrykh et al. 2011). The O_2_ flux ratio of 0:1 in DCMU treated thylakoids was consistent with activity of Reaction 5 (Fig. 1), suggesting peroxidation of lipids and membranes via formation of ^1^O_2_ and/or organic peroxides (Khorobrykh, S. A., Khorobrykh et al. 2011), in the absence of water splitting at PSII. Addition of catalase had no effect on the measured rates (Fig. 4F) confirming that H_2_O_2_ accumulation was not behind the observed O_2_ consumption. This is in line with findings, based on the use of the fluorescent biosensor HyPer2, that PSII cannot generate H_2_O_2_ in the presence of DCMU (Exposito-Rodriguez, Laissue et al. 2017). Increased irradiance also affected the DBMIB and HgCl_2_ treatments. Steady-state O_2_ production by DBMIB treated samples increased by approximately 50%, from 1.0 up to 1.5 μmol O_2_ mg chl^-1^ h^-1^ (Fig. 4C), whilst it increased threefold in the HgCl_2_ samples, from approximately 1.0 to 3.0 μmol O_2_ mg chl^-1^ h^-1^ (Fig. 4D). However, O_2_ consumption rates in both treatments increased by a larger relative amount than O_2_ production, shifting the apparent O_2_ flux ratio towards 1:2 (Fig. 4E). Although, based on the O_2_ flux ratio, this suggested that H_2_O_2_ was accumulating within the thylakoid membrane, we observed that the rate of O_2_ consumption in DBMIB samples was insensitive to catalase (Supplemental Fig. S5 A, we could not measure HgCl_2_ samples with catalase due to the toxicity of Hg^2+^ for the catalase enzyme). We suspect that the isolated thylakoid samples experience strong acceptor side limitation at PSII. This was exacerbated by the higher irradiance which increased the probability of chlorophyll triplet states and therefore ^1^O_2_ formation. The ^1^O_2_ associated lipid/membrane peroxidation was enhanced by the HL treatment, resulting in a strong background rate of O_2_ consumption not associated with electron transport or O_2_^−•^ formation, previously defined as DCMU insensitive O_2_ consumption (Furbank, Badger 1983).

**Figure 4.**
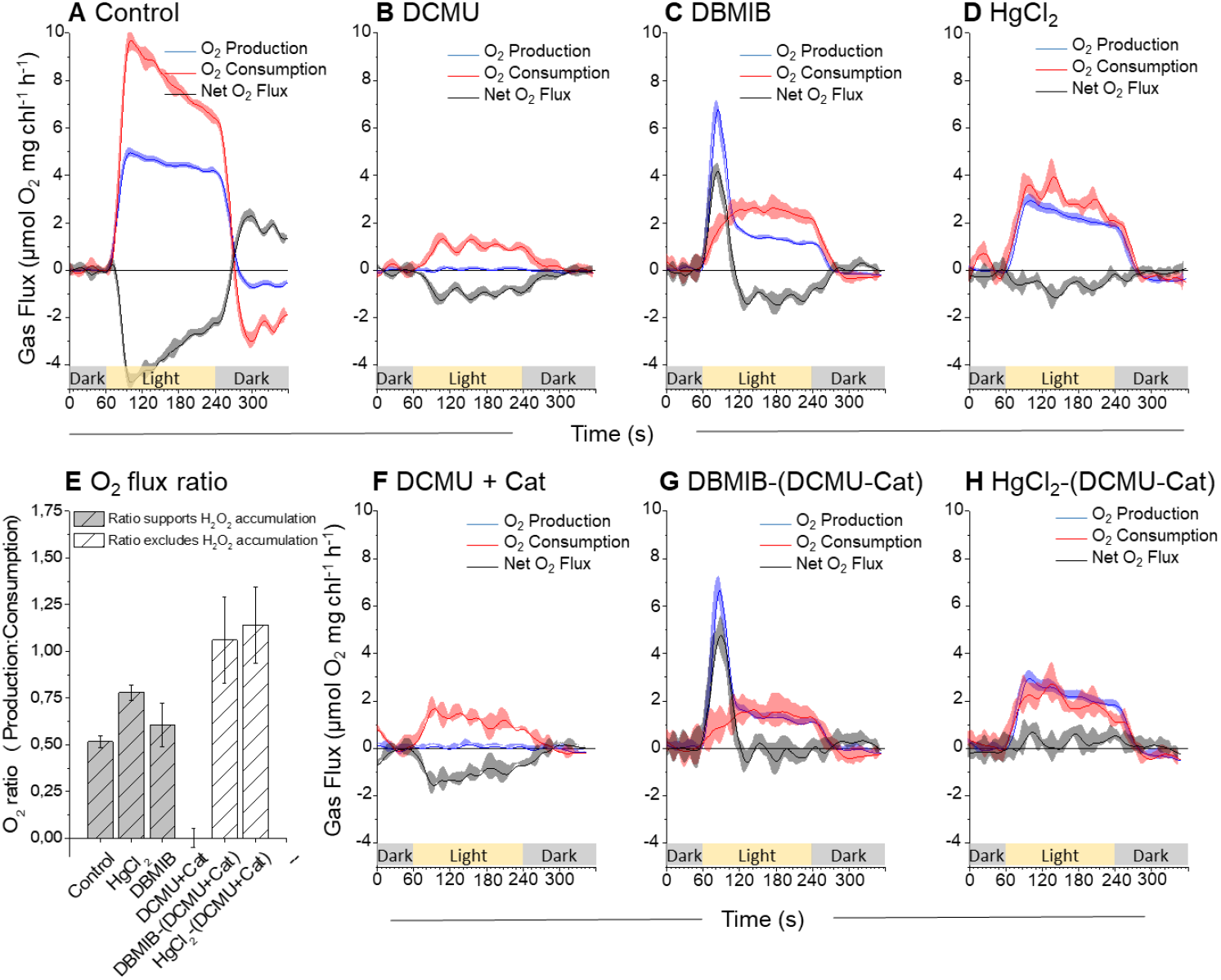
Integrated rates of ^16^O_2_ production and ^18^O_2_ consumption by isolated thylakoid samples, measured simultaneously at 900 μmol photons m^-2^ s^-1^ with MIMS. Illumination of samples represented by yellow bar with grey representing darkness. A, untreated control, B, 50 μM DCMU C, 10 μM DBMIB D, 2 mg ml^-1^ HgCl_2_. O_2_ consumption rates increased more than O_2_ production rates in all inhibited samples, compared to measurements at 120 μmol photons m^-2^ s^-1^. F, 50 μM DCMU + Catalase suggests ^1^O_2_ formation as a likely reason for increased O_2_ consumption, subsequently this curve was subtracted from both DBMIB and HgCl_2_ curves resulting in G, DBMIB-(DCMU + Catalase) and H, HgCl_2_-(DCMU + Catalase). E, Plotting O_2_ production to O_2_ consumption ratios from all curves highlights that subtraction of O_2_ consumption associated with peroxidation of lipids, proteins and membranes by ^1^O_2_ resulted in the return of O_2_ flux ratios that exclude the accumulation of H_2_O_2_. All curves are an average of minimum three representative replicates plotted with standard error. All measurements of isolated thylakoids were performed in measurement buffer containing the uncoupler NH_4_Cl (5mM).

This interpretation was supported when we tested MV+catalase at high light, in which a small catalase insensitive O_2_ consumption was also observed (Supplemental Fig. S5 B). In order to account for the contribution of ^1^O_2_ formation in calculating the absolute O_2_ flux ratios of DBMIB and HgCl_2_ samples, the DCMU + catalase rates were directly subtracted from the DBMIB and HgCl_2_ results (Fig. 4, G and H). Resultant curves from both DBMIB and HgCl_2_ treatments were almost identical to those measured at GL, excluding the slight increase in steady-state gas fluxes. Both exhibited an O_2_ flux ratio of approximately 1:1 (Fig. 4E), further supporting that H_2_O_2_ cannot accumulate within the thylakoid membrane at either PSII, the PQ pool or Cyt-*b*_*6*_*f* complex. This conclusion was further reinforced by the insensitivity to catalase of the HL enhanced O_2_ uptake of DBMIB treated thylakoid samples (Supplemental Fig. S5 A).

### EPR spin trapping data, consistent with conclusions from in-vitro MIMS measurements, supports the function of PTOX

To test conclusions relating to ROS formation in the PETC based on MIMS measurements, we next applied EPR spin trapping to isolated thylakoid samples. We used DIPPMPO, a more lipophilic spin trap with higher sensitivity and greater adduct stability for O_2_^−•^ (DIPPMPO-OOH), OH^−•^ (DIPPMPO-OH), and carbon-centred adducts (DIPPMPO-R) than other commonly used spin traps like DMPO (Villamena 2017). Isolated thylakoids were incubated with DIPPMPO in darkness for five minutes, before sample illumination with actinic light for three minutes at 150 μmol photons m^-2^ s^-1^. Samples were then centrifuged and the supernatant transferred into an EPR capillary for measurement, while the pellet was discarded. To compare the relative capacity for ROS formation at different locations within the PETC compared to untreated controls, the inhibitors DCMU and HgCl_2_ were applied across a series of measurements with MV included as a positive control for enhanced O_2_^−•^ adduct formation. Based on our MIMS results on redox activity of DBMIB (Fig. 3F and Fig. 4, C and G), we excluded DBMIB from EPR measurements as it is also known to alter the redox potential of the Iron-sulphur clusters of PSI and Cyt-*b*_*6*_*f* complex (Malkin 1981), thus possibly producing artefacts and interference in EPR spin trapping of ROS.

To control putative background EPR signals, potentially arising from DIPPMPO incubation with isolated thylakoids and the inhibitors, the EPR response of each sample was measured after five minutes of dark incubation. No EPR signal was produced in darkness (Fig. 5A). Illumination of the spin trap with control thylakoids generated a clear spin trap -ROS adduct signal (Fig. 5B, black trace) which increased 2-3 fold through addition of MV (Fig. 5B, purple trace). We performed a simulation fitting of the experimental EPR spectra to figure out what proportion of different radicals are present in it. Auto simulation was performed by Winsim application using previously reported values of hyperfine coupling constants for DIPPMPO adducts (Chalier, Tordo 2002), setting a 1% limit for g-value and hyperfine coupling constants but enabling automatic adjustment for other parameters during fitting. The best fit simulated spectrum of DIPPMPO radical is obtained (Fig. 5C) for an experimental spectrum of untreated control thylakoids (Fig. 5B, upper trace). It contained approximately 73% O_2_^−•^ adduct (DIPPMPO-OOH), 18 % OH^−•^ (DIPPMPO-OH) and 9% carbon-centred adduct (DIPPMPO-R). Variation of 5 – 10% was observed in the contribution of superoxide and hydroxyl adducts between biological replicates.

**Figure 5.**
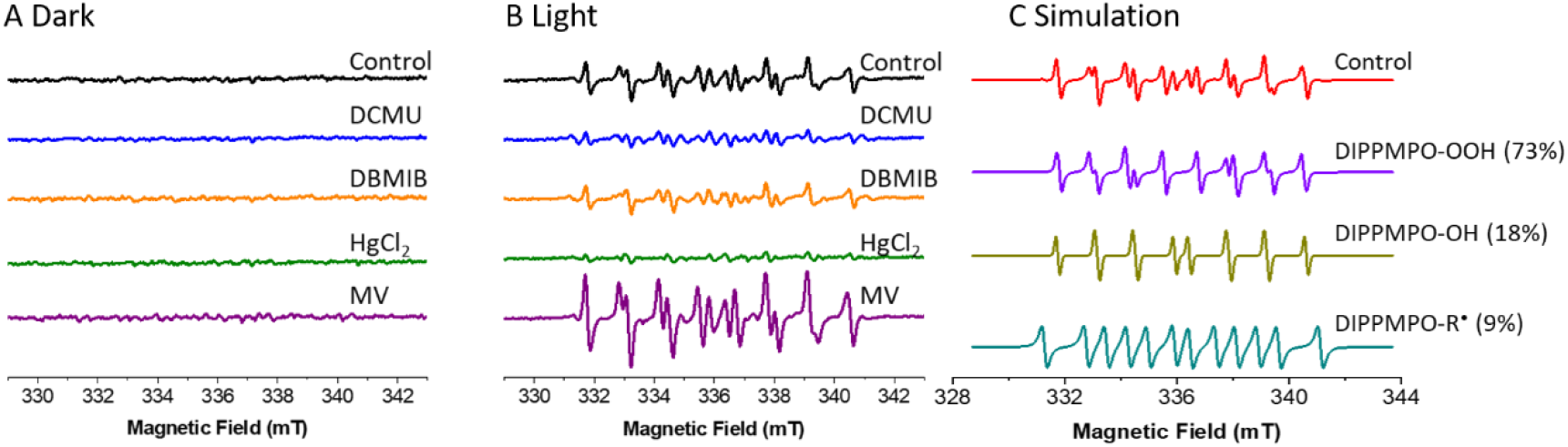
Light induced superoxide formation in isolated thylakoids. Superoxide was measured by spin trapping with DIPPMPO A, in the dark and B, after 3 min of illumination with white light (150 μmol photons m^−2^ s^−1^). Typical spectra of DIPPMPO-OOH with hyperfine splitting constants (*cis a*_P_ 4.968 mT, *a*_N_ 1.314 mT, *a*_H_ 1.102 mT; *trans a*_P_ 4.95 mT, *a*_N_ 1.301 mT, *a*_H_ 1.022 mT), DIPPMPO-OH (*a*_P_ 4.659 mT, *a*_N_ 1.414 mT, *a*_H_ 1.339 mT) and DIPPMPO-R (*a*P 4.59 mT, *a*N 1. 491 mT, *a*H 2.22 mT) adducts were measured in the presence of DCMU (Q_B_–site inhibitor), HgCl_2_ (plastocyanin inhibitor) and MV (Catalyst of O_2_ reduction at PSI). C, simulated spectra of the experimental spectrum of control thylakoids from (B), upper trace consisting of different proportion of each DIPPMPO-OOH, DIPPMP-OH and DIPPMPO-R are shown. Thylakoids were isolated from 6 week old plants, grown under constant light of 120 μmol photons m^−2^ s^−1^ with dark and light cycle of 16/8 h. Thylakoids equivalent to 150 μg chl ml^−1^ were illuminated with actinic light (150 μmol photons m^−2^ s^−1^) in the presence of 50 mM DIPPMPO, 100 μM desferal and 50 mM Hepes– NaOH (pH 7.5) with each electron transfer modulator DCMU (10 μM), HgCl_2_ (2 mg per 150 μg chl) and MV (10 μM). EPR settings were microwave frequency 9.41 GHz, centre field 336.2 mT, field sweep 15 mT, microwave power 5 mW, modulation frequency of 100 kHz, modulation width of 0.05 mT, sweep time 60 s and each EPR spectra were obtained by 5 accumulations of each sample.

The addition of DCMU decreased the measured spin-trap signal by 90% compared to untreated controls (Fig. 5B). A very weak signal in DCMU treated thylakoids comprised approximately 65% carbon centred radicals and only small contributions were observed from O_2_^−•^ and OH^−•^ adducts (Fig. 5B, blue trace). This was consistent with the 0:1 O_2_ flux ratio (Fig. 1, Reaction 5) observed in MIMS data (Fig. 4B) and the peroxidation of lipids and membranes via formation of ^1^O_2_ and/or organic peroxides (Khorobrykh, S. A., Khorobrykh et al. 2011). The addition of HgCl_2_ reduced the adduct signal up to 95% compared to the untreated control samples (Fig. 5B, green trace), despite MIMS data showing that HgCl_2_ only slowed PSII O_2_ evolution, and associated O_2_ consumption, by approximately 40 % (Fig. 3G and 4H). Whilst the O_2_ dynamics strongly suggest that electrons produced through oxidation of water at PSII were accepted by O_2_, the lack of O_2_^−•^ adduct formation infers enzymatic reduction of O_2_ to water, without formation of ROS. Such a pathway is reminiscent of the flavodiiron proteins functional at the PSI acceptor side in lower-order phototrophs and gymnosperms (Ilík, Pavlovič et al. 2017), and makes it conceivable that PTOX performs a similar function in flowering plants. The data is also consistent with function of the theorised PQH_2_/PQH^-^ mechanisms for formation and quenching of O_2_^−•^ (Khorobrykh, Sergey, Tyystjärvi 2018). However, spin-traps are notorious for their inefficiency at penetrating the membranes, making this data difficult to interrogate further. Despite this, the suppression of the EPR spin-trap signal (Fig. 5B), in conjunction with the 1:1 O_2_ flux ratio measured with MIMS (Fig. 3G and 4H), rules out the accumulation of H_2_O_2_ in HgCl_2_ treated thylakoid samples. As HgCl_2_ enabled the reduction of all components upstream of PSI, these data offer robust evidence that stable accumulation of H_2_O_2_ requires PSI activity.

### Detecting photosynthetic H_2_O_2_ accumulation via O_2_ stoichiometry in leaf discs

To test the applicability of our in-vitro findings we developed a MIMS methodology to estimate O_2_ flux ratios in-vivo (Arabidopsis leaf discs), under similar conditions to those tested with isolated thylakoids. As the primary terminal electron acceptor in leaves is CO_2_ and mitochondrial respiration is a large contributor to O_2_ consumption, it was not possible to simply divide the gross O_2_ production rate by the gross O_2_ consumption rate. Alternatively, we estimated the O_2_ flux ratio associated specifically with the steady-state WWC determined in the absence of photorespiration. Assuming a 1:1 ratio between ^16^O_2_ evolution and ^13^CO_2_ (artificially enriched) fixation by Rubisco during oxygenic photosynthesis, we determined the difference between the production of electrons at PSII and the consumption of electrons by the Calvin-Benson-Bassham (CBB) cycle. This difference provided an upper limit of ^16^O_2_ production specifically linked to the WWC. The rate of ^18^O_2_ consumption associated with the Mehler reaction was then determined by subtracting the rate of ^12^CO_2_ produced by mitochondrial respiration, from the gross rate of ^18^O_2_ consumption (assuming a 1:1 ratio between CO_2_ production and O_2_ consumption during aerobic mitochondrial respiration). By simultaneously measuring the fluxes of four stable isotopes, representing isotopologues of both O_2_ and CO_2_, we could estimate the steady-state rate of O_2_ production and O_2_ consumption associated specifically with the WWC and hence, estimate the in-vitro and in-vivo O_2_ flux ratio according to the scheme in Figure 1.

Arabidopsis leaf discs (12.5 mm in diameter) were floated overnight in near darkness on water containing 10 μM DCMU, 10 μM MV or 10 μM DBMIB (or plain water – untreated control), then sealed into the MIMS cuvette and purged with air scrubbed of all ^12^CO_2_ (Soda Lime, Li-Cor USA). To this 3% ^13^CO_2_ and 2% ^18^O_2_ were injected by volume (total O_2_ approximately 21%). After a few minutes darkness to ensure isotopic equilibrium throughout the sample, data acquisition was initiated. An illumination protocol of three minutes darkness, five minutes at GL, five minutes at HL and three minutes darkness was applied. In the data, all samples exhibited an apparent ‘peak’ of ^18^O_2_ production during transition from GL to HL, and ^18^O_2_ uptake following transition from HL to darkness. This trend was also observed in the blank control measurements (Supplemental Fig. S6) indicating that it was an artefact, potentially a light effect on the membrane. To avoid this artefact and emphasise the steady-state results in the main panels, the integrated gas flux rates versus time (Fig. 6, A, B, D and E) are displayed with breaks in the x-axis. For reference, the inset figures display the full curves. All rates used to calculate the O_2_ flux ratios (Fig. 6F) are based on the averaged steady-state fluxes measured from minimum three biological replicates. These values are presented as a table in the supplementary data (Supplemental Tab. S1).

**Figure 6.**
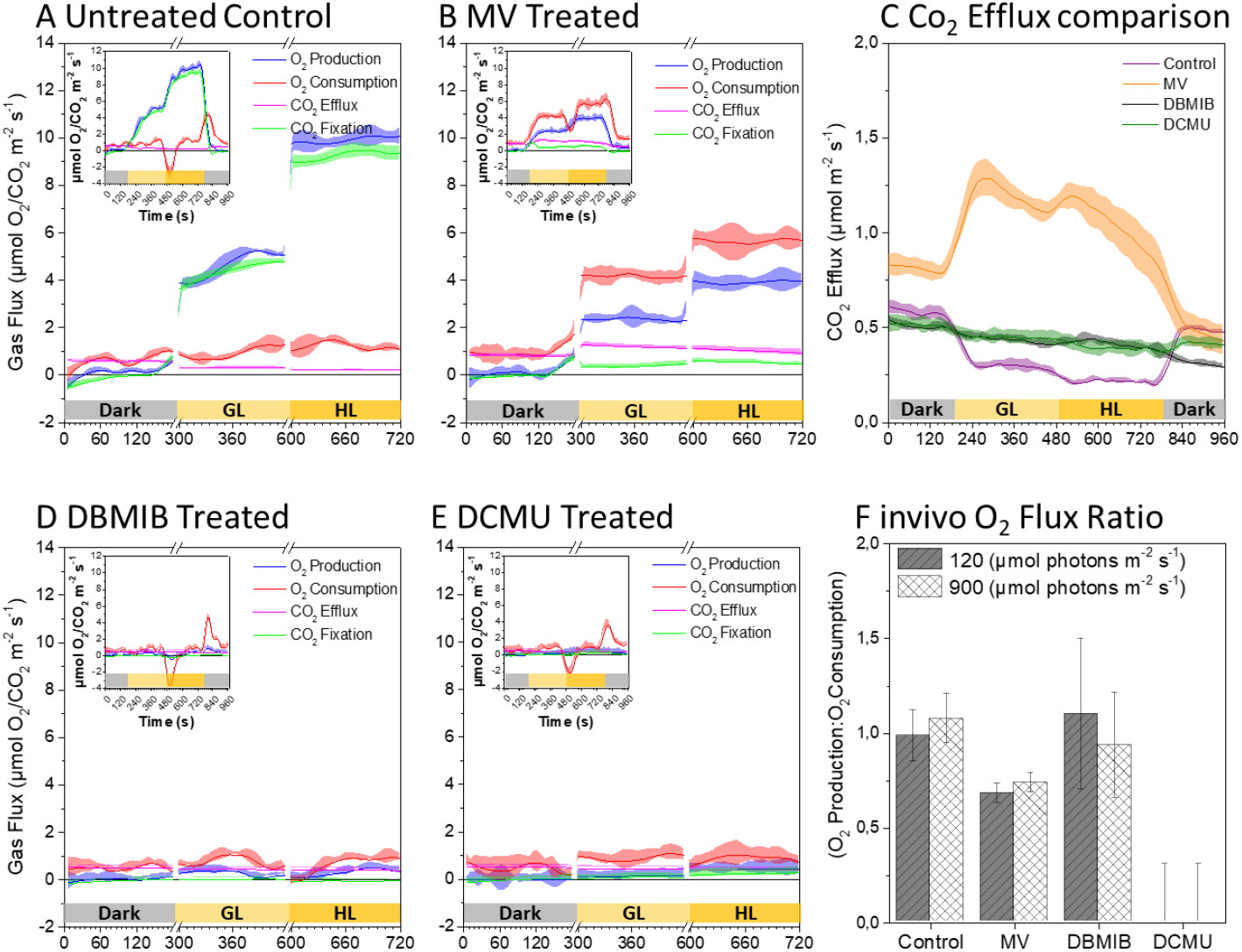
The integrated rates versus time of O_2_ and CO_2_ fluxes from intact leaf discs measured at three different light conditions. Rate versus time plots of A, control, B, MV, D, DBMIB, E, DCMU infiltrated leaf discs. Inset shows complete curves and the main figure highlights steady-state rates across the dark and two light intensities. GL = 120 and HL = 900 μmol photons m^-2^ s^-1^ respectively. For direct comparison, C, shows CO_2_ efflux rates (mitochondrial respiration) from the four treatments. F, O_2_ flux ratio associated with Mehler reaction calculated as described in text. All curves average of minimum three representative replicates plotted with standard error. The apparent mismatch between activation of light and photosynthetic activity is a result of the integration of rates over time required to minimise noise and allowing a focus on steady state rates.

Untreated control leaf discs (Fig. 6A) performed as expected in a high CO_2_ environment. Rates of O_2_ production and CO_2_ fixation (Fig. 6, blue and green curves) were strongly correlated across all irradiances. The ^12^CO_2_ efflux generated by mitochondrial respiration apparently decreased in correlation with increasing irradiance (Fig. 6, A-E, pink line, Fig. 6C, highlighted by purple line). This result was likely a product of CO_2_ re-fixation (Busch, Sage et al. 2013), as such dark respiration was assumed to be constant when calculating the in-vivo O_2_ flux ratio from untreated control discs, as concluded by (Farquhar, Busch 2017). The rate of O_2_ production providing electrons used specifically in the Mehler reaction of control discs was estimated at 0.4 ± 0.3 and 0.7 ± 0.6 μmol O_2_ m^-2^ s^-1^ at GL and HL respectively. This matched the estimated rate of Mehler O_2_ consumption of 0.4 ± 0.1 and 0.6 ± 0.4 μmol O_2_ m^- 2^ s^-1^ at GL and HL.

The resulting in-vivo O_2_ flux ratio of approximately 1.0 (GL = 1.0 ± 0.1, HL = 1.1 ± 0.1) discounted accumulation of H_2_O_2_, as expected with an elevated CO_2_ atmosphere. Despite differences in methodology, these rates compare well with previous MIMS based estimates of Mehler specific O_2_ consumption under similar conditions (Mubarakshina, Ivanov 2010).

The DBMIB treatment completely impaired ^13^CO_2_ fixation (Fig. 6D, green line) and had no effect on ^12^CO_2_ efflux (Fig. 6C, black line). However, light dependent O_2_ production was evident and it correlated with an increase in the rate of O_2_ consumption above that required for mitochondrial respiration. This result supports the function of an O_2_ reduction pathway within the thylakoid membrane, observed in-vitro, and the calculated O_2_ flux ratio of approximately 1.0 (GL = 1.1 ± 0.4 and HL = 0.9 ± 0.3) reinforced the in-vitro conclusions that H_2_O_2_ cannot accumulate within the reduced PQ pool. The DBMIB dependent O_2_ production ‘peak’ was not observed in the leaf discs, potentially due to chlororespiration during dark incubation preceding the measurements that may have fully reduced all DBMIB in the samples before illumination. Contrasting the earlier experiments, DCMU infiltration was not completely effective, although it severely impaired O_2_ production during illumination (Fig. 6E). However, this O_2_ production was matched by commensurate CO_2_ fixation during illumination (GL = 0.1 ± 0.06, HL = 0.2 ± 0.2 μmol m^-2^ s^-1^). Significantly, the illuminated O_2_ consumption rate doubled from a dark rate of 0.4 μmol O_2_ m^-2^ s^-1^ to approximately 0.8 μmol O_2_ m^-2^ s^-1^. In the absence of any change to the respiratory CO_2_ efflux (Fig. 6C, green trace) and with all PSII activity accounted for by CO_2_ fixation, the light dependent O_2_ consumption was most likely a product of ^1^O_2_ formation and associated peroxidation of lipids, proteins and membranes. The calculated O_2_ flux ratio of zero (Fig. 6F) was in-line with our in-vitro MIMS and EPR results and in agreement with published literature that H_2_O_2_ cannot accumulate at PSII in the presence of DCMU (Exposito-Rodriguez, Laissue et al. 2017).

As a final positive control to test the methodology in estimating H_2_O_2_ accumulation, we infiltrated leaf discs with MV. In leaf discs MV competes with CO_2_ fixation reactions for reductant at PSI, impairing rates of CO_2_ fixation compared to untreated control discs. The accumulation of ATP that would otherwise be used in the CBB cycle limits the supply of phosphate for further ATP production. The subsequent impairment of ATP-synthase results in the accumulation of a strong proton gradient across the thylakoid membrane, which impairs O_2_ production and likely results in a counterintuitively reduced PQ pool as previously described (Shapiguzov, Nikkanen et al. 2020). The illuminated rate of O_2_ consumption in MV treated leaf discs was approximately twofold the rate of O_2_ production, suggesting the accumulation of H_2_O_2_ (Fig. 6B). However, in darkness MV approximately doubled the ^12^CO_2_ efflux rate associated with mitochondrial respiration compared with all other samples (Fig. 6C, orange line), as observed by (Scarpeci, Valle 2008). Uniquely for the MV treated samples, the ^12^CO_2_ efflux rate was further enhanced by illumination. Assuming this was due to mitochondrial respiration with a 1:1 respiratory quotient accounted for a significant portion of the increase in O_2_ consumption. This observation was only possible due to our application of the ^13^CO_2_ offset method, which enabled an accurate O_2_ flux ratio of 0.7 ± 0.05 to be determined at both GL and HL. This ratio supports the formation and accumulation of some photosynthetically derived H_2_O_2_, as more O_2_ was consumed than produced by PSII. However, the ratio also suggests that ROS scavenging components associated with the WWC were able to manage approximately half of the H_2_O_2_ produced in the presence of MV, at the impaired rates of PSII activity. Although the signal to noise ratio of this in-vivo method could not discriminate subtle events, the in-vivo data broadly supported the in-vitro conclusions that 1) H_2_O_2_ could not accumulate at PSII 2) an O_2_ reduction pathway operates in the presence of DBMIB which precludes the accumulation of H_2_O_2_. In addition, application of ^13^CO_2_ to discriminate mitochondrial respiration from CO_2_ fixation by the CBB cycle unexpectedly revealed a light dependent stimulation of mitochondrial respiration in MV treated leaf discs, which we examined further.

### Functional evidence suggests that H_2_O_2_ accumulation induces cooperation between chloroplasts and mitochondria

Infiltration with MV resulted in a light dependent increase in mitochondrial ^12^CO_2_ efflux. This respiratory ‘burst’ occurred after the light dependent increase in ^18^O_2_ consumption (Fig. 6B, compare the red and pink curves, note that apparent ‘early’ light response in data is an artefact from integrating data points over 30 seconds, all gas fluxes were integrated equally). To further probe this result we vacuum infiltrated leaves with catalase and with rotenone (inhibitor of mitochondrial Complex-I). Neither compound affected the gas exchange trends of ^12^CO_2_ efflux in comparison with untreated control (infiltrated with water) discs (Supplemental Fig. S7, compare blue, pink and black curves). When leaves were infiltrated with MV + rotenone, samples exhibited a similar light induced respiratory burst of ^12^CO_2_ efflux as the standard MV infiltrated leaves (Fig. 6C, compare the orange curve to the red curve in Supplemental Fig. S7). However, leaves infiltrated with MV + catalase shared dynamics with DCMU and DBMIB infiltrated discs, showing no light dependent burst of ^12^CO_2_ efflux (Fig. 6C, compare green and black curves with the green curve of Supplemental Fig. S7). Therefore, the light dependent ^12^CO_2_ burst was only apparent following infiltration with MV, was not affected by rotenone but was blocked by catalase. We speculate this may be evidence of H_2_O_2_ signaling to the mitochondria (Cui, Brosché et al. 2019), which could relate to ‘malate cycling’ (Zhao, Yu et al. 2020), possibly evidence that the mitochondria are primed to process an influx of malate during a prolonged PSI acceptor limitation, providing a buffer to minimise ROS accumulation and damage at PSI (Noguchi, Yoshida 2008). As no malate was formed in leaves infiltrated with MV, the primed mitochondria increased the rate of decarboxylation reactions due to an increased sink availability.

## Discussion

ROS produced photosynthetically in the chloroplast’s thylakoid membrane play a key role in photodamage, environmental sensing and photosynthetic regulation. Within this paradigm it is proposed that relatively stable ROS, such as H_2_O_2_, can be exported from the chloroplast to act directly as a retrograde signal in fine-tuning nuclear gene expression during plant acclimation to changing environments (Gollan, Aro 2020, Exposito-Rodriguez, Laissue et al. 2017), or as a trigger for cellular processes such as stomatal closure (Wang, He et al. 2016, Iwai, Ogata et al. 2019). Models of ROS signaling require that the sources of, and responses to (signaling pathway) specific ROS are well understood. Whilst chloroplast H_2_O_2_ signaling pathways are slowly being defined (Gollan, Aro 2020, Crisp, Ganguly et al. 2017, Bechtold, Richard et al. 2008, Rossel, Wilson et al. 2007, Dietz, Mittler et al. 2016), the endogenous sources of specific ROS in the thylakoid membrane and the environmental conditions that lead to their formation are still subject to debate. For example, the current literature supports the generation of a stable H_2_O_2_ pool, capable of export from the chloroplast, in multiple sites of PETC including the reduction of O_2_ at PSII (Khorobrykh, Andrey 2019, Tiwari, Pospisil 2009), the PQ pool (Khorobrykh, S. A., Ivanov 2002, Mubarakshina, Ivanov 2010, Khorobrykh, Sergey A., Karonen et al. 2015), PTOX (Heyno, Gross et al. 2009), Cyt-*b*_*6*_*f* complex (Baniulis, Hasan et al. 2013) and PSI (Mehler 1951, Kozuleva, Ivanov et al. 2020). This broad range of candidates complicates models of ROS signaling, resulting in debates like those between Wang et al (Wang, He et al. 2016) and Iwai et al (Iwai, Ogata et al. 2019). Whilst both agree that stomatal closure is triggered by chloroplast derived H_2_O_2_, the former argues in favour of reduced PQ as the source H_2_O_2_ and the latter concludes that it must be sourced from PSI. These discrepancies, together with our recent finding that the inhibitor DNP-INT, used historically in this field, fails to completely block re-reduction of P700^+^ (Fitzpatrick, Aro et al. 2020) prompted us to re-evaluate the ROS formation by specific PETC components of the thylakoid membrane.. As the application of DNP-INT in previous studies (Borisova-Mubarakshina, M. M., Naydov et al. 2018, Khorobrykh, S. A., Ivanov 2002) likely failed to fully inhibit the PSI-Mehler reaction, artefacts may have been reported which could complicate the models of ROS signaling. By thoroughly testing the efficacy of all chosen inhibitors with P700 spectroscopy, and directly quantifying the simultaneous O_2_ production and consumption reactions with MIMS, we have taken here a new approach to measuring site-specific ROS formation within the PETC, and particularly the formation of a steady pool of H_2_O_2_, which minimises the number of assumptions to be made during analysis.

### H_2_O_2_ accumulation via O_2_ photoreduction in isolated thylakoids occurs exclusively at PSI

Measuring the stoichiometry of O_2_ produced and consumed by isolated thylakoids during illumination enabled the calculation of an O_2_ flux ratio, which was anticipated to reach 1:2 when H_2_O_2_ accumulated as described previously by (Allen 1977, Asada 1999, Asada 2006) (Fig. 1). As expected, we observed a 1:2 O_2_ flux ratio in untreated control and MV treated thylakoid samples (Fig. 3H and 4E). Further addition of catalase to these thylakoid samples pushed the ratio back to 1:1 (Fig. 3, B and D), confirming that accumulation of H_2_O_2_ was responsible for the observed 1:2 O_2_ flux ratio. This result fully supports the known fact that the PSI-Mehler reaction generates a stable pool of H_2_O_2_ (Mehler 1951). In stark contrast to O_2_ flux results with control and MV treated thylakoids, the isolated thylakoids incubated with HgCl_2_ and DBMIB, which we demonstrated with concomitant measurements of P700 redox kinetics to completely block all PSI function (Fig. 2, A and B), exhibited 1:1 O_2_ flux ratios. These stoichiometric O_2_ production and consumption experiments undoubtedly demonstrate that efficient inter-chain PETC inhibitors completely block the accumulation of H_2_O_2_ and, conversely, a stable pool of H_2_O_2_ can be acquired in isolated thylakoids only via the activity of PSI.

### Separating the *formation* of H_2_O_2_ from its *accumulation* within the thylakoid membrane

Stable accumulation of H_2_O_2_ produced a 1:2 O_2_ flux ratio in our experiments, being consistent to support the function of a putative direct H_2_O_2_ retrograde signal. In contrast, the 1:1 O_2_ flux ratio that implies no stable accumulation of H_2_O_2_ for export, questions the involvement of thylakoid components upstream of PSI in the H_2_O_2_ related long distance regulatory processes. However, a significant body of previous research suggests that H_2_O_2_ can form upstream of PSI (the PQ pool (Khorobrykh, S. A., Ivanov 2002, Mubarakshina, Ivanov 2010, Khorobrykh, Sergey A., Karonen et al. 2015), PTOX (Heyno, Gross et al. 2009), Cyt-*b*_*6*_*f* complex (Baniulis, Hasan et al. 2013)). Whilst our data seems to contradict these works, we postulate that the discrepancies merely highlight a difference in our approach, which relies on the strength of the MIMS data in separating H_2_O_2_ *formation* from H_2_O_2_ *accumulation*. Recall that illumination of control and MV treated thylakoid samples generated significant quantities of H_2_O_2_ (Fig. 3, A and C), yet a 1:1 O_2_ flux ratio was observed in the presence of catalase (Fig. 3, B and D). In the same manner, it is plausible that H_2_O_2_ may form within the thylakoid membrane upstream of PSI as previously suggested (Mubarakshina, Ivanov 2010, Khorobrykh, S. A., Ivanov 2002, Khorobrykh, Sergey A., Karonen et al. 2015, Heyno, Gross et al. 2009, Baniulis, Hasan et al. 2013). Nevertheless, the MIMS data unequivocally shows that such H_2_O_2_ is not stable and must be rapidly broken down, analogous to the catalase treated control samples. Importantly, PSII oxygen evolution was active in the presence of DBMIB and HgCl_2_ (Fig. 3, B and D), making it plausible that O_2−•_ was formed and therefore also H_2_O_2_ was apparently being transiently produced in the absence of PSI. It is conceivable that such unstable H_2_O_2_ may be positively detected by sensitive dyes (Cathcart, Schwiers et al. 1983), spin-traps, specific sensor proteins (Villamena, Zweier 2004) or other sensitive methods of H_2_O_2_ detection. For this reason, the capacity of the MIMS method to discriminate between H_2_O_2_ that accumulated, versus H_2_O_2_ that briefly formed, is a functionally significant distinction from the perspective of the putative H_2_O_2_ retrograde signal.

### Pathway(s) of O_2_ photoreduction within the thylakoid membrane

MIMS measurements of thylakoid samples incubated with HgCl_2_ (Fig. 3G and 4H) and both thylakoid and leaf samples incubated with DBMIB (Fig. 3F, 4G and 6D) maintained steady-state O_2_ production during illumination. This PSII activity was associated with commensurate O_2_ consumption resulting in a 1:1 O_2_ flux ratio. Both DBMIB and HgCl_2_ treated thylakoid samples exhibited similar O_2_ flux rates at GL which did not increase during HL illumination in the DBMIB treated samples, a dynamic also observed in DBMIB infiltrated leaf discs (Fig. 6D and Supplemental Tab. S1,). Conversely, the O_2_ flux rate doubled in response to HL in the HgCl_2_ thylakoid samples (Supplemental Fig. S1, direct comparison of thylakoid samples). As the PSII contribution to O_2−•_ formation is relatively low (Khorobrykh, Andrey 2019), the DBMIB pathway (excluding Cyt-*b*_*6*_*f*) tested the maximum rate of O_2_ reduction resulting from direct interaction between reduced Q_A_, Q_B_ or the PQ pool and O_2_, potentially via the theorised PQH_2_/PQH^-^ O_2_^−•^ formation and quenching mechanism (Khorobrykh, Sergey, Tyystjärvi 2018) and/or directly via PTOX (Cournac, Josse et al. 2000). That the rate was saturated at a low irradiance was consistent with in-vitro characterisations of the PTOX pathway (Nawrocki, Tourasse et al. 2015). In comparison, the reduction of PQ pool and Cyt-*b*_*6*_*f* in HgCl_2_ treated thylakoids correlated to an increased rate of both O_2_ production and consumption at HL, which supports the widely held hypothesis that Cyt-*b*_*6*_*f* can generate O_2_^−•^ as demonstrated by (Baniulis, Hasan et al. 2013) (as HgCl_2_ was toxic for leaf discs, a direct comparison between in-vitro and in-vivo HgCl_2_ samples was impossible). We unexpectedly found that application of the specific PTOX inhibitors n-propyl- and octyl-gallate increased the rates of both O_2_ production and consumption (Supplemental Fig. S4), confounding our attempt to examine the specific contribution of PTOX to the rate of O_2_ reduction. The absence of an EPR spin-trap signal in HgCl_2_ treated thylakoids supports a route of catalysed O_2_ reduction within the thylakoid membrane, possibly at PTOX (Fig. 5B, green curve). These results suggest that (i) thylakoid membranes apparently support at least two separate routes of O_2_ reduction upstream of PSI, (ii) both routes of O_2_ reduction exhibit a lower absolute capacity than the PSI-Mehler reaction and (iii) none of the O_2_ reduction pathways upstream of PSI produce a stable pool of H_2_O_2_ that could be exported as a retrograde signal.

### A simpler model for H_2_O_2_ signaling

Our conclusions broadly support what is already described regarding the *formation* of ROS by photosynthetic processes in the chloroplast (Kozuleva, Ivanov et al. 2020), including our observation of multiple distinct pathways for ROS formation associated with the thylakoid membrane. We have not ruled out that O_2_^−•^ and possibly H_2_O_2_ form upstream of PSI and can be detected by rapidly reacting dyes (Cathcart, Schwiers et al. 1983), spin-traps or specific sensor proteins (Villamena, Zweier 2004). Despite this, our results dismiss the possibility that any stable pool of H_2_O_2_ can *accumulate* upstream of PSI. This finding has significant consequences for models of chloroplast H_2_O_2_ retrograde signaling, as it implies that any H_2_O_2_ exported from the chloroplast was formed at PSI. In the absence of added Fd, isolated thylakoid samples do not exhibit strong rates of O_2_ reduction (Furbank, Badger 1983). This implicates the acceptor side of PSI in O_2_^−•^ formation. In addition, activity of O_2_ reduction was efficiently quenched by the addition of NADP^+^ (Furbank, Badger 1983), suggesting that the acceptor side capacity of PSI regulates the rate of O_2_^−•^ formation. As no other potential source of H_2_O_2_ formation could generate a stable H_2_O_2_ pool within the thylakoid membrane during our experiments, the data suggest that only the PSI acceptor side contributes to H_2_O_2_ involved in direct H_2_O_2_ retrograde signaling (Gollan, Aro 2020, Exposito-Rodriguez, Laissue et al. 2017). Therefore, the export of H_2_O_2_ from the chloroplast specifically communicates PSI acceptor limitation to the broader cell.

Based on the available data we propose that H_2_O_2_ sufficiently accumulates for export from chloroplasts only under conditions where PSI acceptors are fully reduced, and the antioxidant pathways of the WWC have become overwhelmed. This is in line with the absence of catalase (which does not require reductant to function or exhibits H_2_O_2_ sensitivity (Mhamdi, Queval et al. 2010)) from chloroplasts, which instead rely on pathways of H_2_O_2_ detoxification that are sensitive to excess H_2_O_2_ accumulation (Kitajima, Nii et al. 2010). In this model, the peroxiredoxin, ascorbate and ascorbate peroxidase enzymes of the Mehler WWC act as a buffer, decreasing the sensitivity of any chloroplast derived H_2_O_2_ signal to short term and transitory stress events. Therefore any H_2_O_2_ retrograde signal, which is the quickest to trigger a regulatory response during HL stress (Gollan, Aro 2020), only forms once environmental conditions potentially leading to PSI damage start to dominate (Tiwari, Mamedov et al. 2016). Such a mechanism avoids nuclear responses during transient stresses, such as sun flecks or patchy cloud cover. This avoids potentially premature down-regulation of key photosynthetic processes such as light harvesting (Borisova-Mubarakshina, Maria, Ivanov et al. 2015) during otherwise low-light conditions. Whilst short term stress events can be mitigated via the enzymes of the WWC and other regulatory processes such as non-photochemical quenching (NPQ) and photosynthetic control at Cyt-*b*_*6*_*f* (Shimakawa, Miyake 2018), longer term environmental shifts that lead to chronic over-reduction of the PSI acceptor pool will result in the deactivation of the WWC’s antioxidant enzymes, followed by accumulation and eventual export of H_2_O_2_. In this model the PSI acceptor side specific H_2_O_2_ retrograde signal is a mechanism to convey information from chloroplasts to the nucleus when dominant environmental conditions change and the PSI acceptor side faces chronic limitation. This provides the nucleus with a mechanism to efficiently respond for acclimation to a new environment, whilst avoiding nuclear responses to short-term perturbations such as sun flecks.

### Possible evidence of direct H_2_O_2_ triggered cooperation between chloroplast and mitochondria

In order to estimate the O_2_ flux ratio associated specifically with the Mehler reaction of leaf discs, we artificially enriched the leaf disc sample atmosphere to 2% with ^13^CO_2_. This enabled us to discriminate between photosynthetic assimilation of ^13^CO_2_ and the efflux of respiratory ^12^CO_2_ (Fig. 6). With this method we observed that leaf discs infiltrated with MV exhibited a sharp increase in ^12^CO_2_ efflux from mitochondrial respiration during illumination. The light dependant burst of ^12^CO_2_ efflux was absent in leaves infiltrated with DCMU or DBMIB, excluding oxidation of the PSI acceptor side as a cause. The ^12^CO_2_ burst was maintained in leaves infiltrated with MV + Rotenone, yet inhibited when leaves were infiltrated with both MV+catalase. These results suggest that H_2_O_2_ formed by MV at the acceptor side of PSI may have triggered an increase in mitochondrial respiration. This fits with a growing body of evidence that mitochondria and chloroplasts interact to optimise the photosynthetic performance. From an efficiency perspective, the organelles co-locate to improve C_3_ photosynthesis by maximising the reassimilation of respired CO_2_ (Busch, Sage et al. 2013). From a regulatory perspective it has long been known that mitochondria can accept excess electrons from the chloroplast’s thylakoid membrane in leaves (Noguchi, Yoshida 2008) and it was recently confirmed that mitochondrial AOX expression reduces ROS damage, and was necessary for HL acclimation in green algae (Kaye, Huang et al. 2019). The term ‘malate circulation’ has been proposed (Zhao, Yu et al. 2020) to describe the various regulatory functions spanning both organelles, potentially including programmed cell death, that relate to the ‘malate valve’ (Selinski, Scheibe 2019).

To explain our observation that MV infiltration increased the CO_2_ efflux measured from leaf discs during illumination in a catalase sensitive manner, we propose that H_2_O_2_ produced by MV was exported from the chloroplast, which then triggered upregulation or ‘priming’ of an oxidative pathway in surrounding mitochondria, potentially AOX (based on work with *Chlamydomonas* (Kaye, Huang et al. 2019)). However, in our experiments, where malate was not exported from the chloroplast and the mitochondria were primed by artificially produced H_2_O_2_, the upregulated pathway increased the oxidative sink capacity within the mitochondria. To satisfy this increased sink, the rate of decarboxylation reactions may have been upregulated, an effect already observed in mitochondria in the absence of photosynthesis (Scarpeci, Valle 2008).

It is conceivable that the pre-incubation with MV contributed to priming this response, hence its rapid activation during illumination, although the speed of the response may be evidence that this is a critical pathway to facilitate protection of PSI during acute acceptor limitation. It precedes down-regulation of photosynthetic processes triggered by H_2_O_2_ retrograde signaling, which take at least half an hour (Gollan, Aro 2020), during which time the interaction may function to provide the PSI acceptor side with either NADP^+^ via the malate valve (Noguchi, Yoshida 2008), or CO_2_ via the upregulated carboxylation reactions observed during our measurements. The latter response could be of great significance if the stomata also close due to a H_2_O_2_ trigger (Iwai, Ogata et al. 2019). This explanation fits our proposed model of PSI specific H_2_O_2_ signaling. It suggests that H_2_O_2_ export from the chloroplast simultaneously signals down regulation of photosynthetic processes during prolonged acceptor limitation, and upregulation of a pathway to temporarily increase the PSI acceptor capacity via the malate valve and potentially directly via the increased availability of respired CO_2_.

### Concluding remarks

We have demonstrated that a stable pool of H_2_O_2_ can accumulate in the thylakoid membrane only via the PSI-Mehler reaction. This explicitly excludes the involvement of other PETC components in putative chloroplastic H_2_O_2_ signaling pathways, with significant implications for models of H_2_O_2_ involvement in processes like stomatal regulation or retrograde signaling. We have confirmed the function of separate O_2_ consumption pathways within the thylakoid membrane, involving the PQ pool and Cyt-b_6_f, although neither demonstrated the capacity to accumulate H_2_O_2_. We propose a model for chloroplastic H_2_O_2_ retrograde signaling based on the PSI acceptor side capacity. This enables efficient nuclear regulation in response to chronic environmental changes, whilst avoiding unnecessary responses to short term stress caused by conditions such as sun-flecks or patchy cloud cover. Finally, we have discovered potential evidence for H_2_O_2_ triggered cooperation between the chloroplast and mitochondria. We suggest that this may alleviate PSI acceptor side stress when the antioxidant systems of the WWC are overwhelmed and PSI is more susceptible to damage whilst nuclear regulation, activated by H_2_O_2_ retrograde signaling, takes effect.

## Methods

### Plant Material and thylakoids isolation

The *Arabidopsis thaliana* plants were grown at atmospheric CO_2_ under dark/light cycle of 16/8 h at 120 μmol photons m^-2^ s^-1^. Thylakoids were isolated from 6 weeks old plants using standard method as described earlier (Tiwari, Mamedov et al. 2016). All measurements were performed with freshly isolated thylakoids in measurement buffer containing: 330 mM Sorbitol, 5 mM MgCl_2_, 10 mM NaCl, 5 mM NH_4_Cl, 50 mM Hepes pH 7.6, unless stated otherwise. Measurements on intact leaf or leaf discs on Dual Pam and MIMS were performed using the same plants. For infiltration of electron transfer inhibitors/modulators in leaves, detached leaves were floated at 25 °C in darkness for 1 h in either H_2_O (control) or H_2_O + 10 μM DBMIB or H_2_O + 10 μM DNP-INT or H_2_O + 10 μM MV.

### P700 redox kinetics measurements

Detached leaves and isolated thylakoids were used for P700 redox kinetics measurements using Dual-Pam-100 (Walz, Germany). The P700 was oxidised under continuous far red (FR) light. The partial and complete re-reduction of P700 by electrons flow from PSII was achieved by shooting two short pulses of saturating actinic light pulses *i*.*e*. single turnover (ST) 50μs and multiple turnover (MT) 50ms, over the FR light oxidised P700 (Tiwari, Mamedov et al. 2016).

### Superoxide measurement

Spin trapping for O_2_^−•^ was performed in isolated thylakoids using Miniscope (MS5000) Electron Paramagnetic Resonance (EPR)-spectrometer equipped with variable temperature controller (TC-HO4) and Hamamatsu light source (LC8). The isolated thylakoids equivalent to 150 μg ml-1 Chl were illuminated under actinic light (150 μmol photons m^-2^ s-^1^) for 180 s in the presence of 5-(Diisopropoxyphosphoryl)-5-methyl-1-pyrroline-N-oxide 2-Diisopropylphosphono-2-methyl-3,4-dihydro-2H-pyrrole-1-oxide (DIPPMPO) spin trap (50 mM) in 50 mM Hepes-NaoH (pH 7.5) with 50 μM desferol. Subsequently, the samples were centrifuged at 6500×g for 5 min and supernatant was used for EPR measurements. The electron transport inhibitors DCMU (10 μM), DBMIB (10 μM), DNP-INT (10 μM), HgCl2 (2 mg ml^-1^) and MV (10 μM) were added in reaction medium when indicated prior to the measurements. The measurements were conducted at frequency 9.41 GHz, centre field 3363 G, field sweep 150 G, microwave power 3 mW and modulation frequency of 100 kHz with modulation width of 2G. The final spectra were obtained by 5 accumulations of each sample of minimum 3 to 5 biological replicates.

### Gas exchange measurements in liquid samples

The MIMS liquid cuvette consisted of an in-house modified Hansatech O_2_ electrode body, with the silver/platinum electrode replaced by a stainless-steel assembly supporting a Teflon membrane (Hansatech, UK) attached directly to the High Vacuum inlet of a Thermo Sentinel-PRO magnetic sector mass spectrometer (Thermo-Fisher USA), collecting masses 32 and 36 with a total cycle time of approximately 4.5 seconds. Freshly isolated thylakoid were stored on ice in darkness. For each run sufficient measurement buffer (containing: 330 mM Sorbitol, 5 mM MgCl_2_, 10 mM NaCl, 5 mM NH_4_Cl, 50 mM Hepes pH 7.6) was loaded into the cuvette with the addition, via syringe (approximately 50 μg ml^-1^ chlorophyll) to a final volume of 1000 μl. In darkness the sample was purged with N_2_ to minimise background ^16^O_2_ before a bubble of ^18^O_2_ (99% Cambridge Isotope Laboratories Inc, UK) was loaded into the stirring liquid, bringing the concentration of the heavier isotope up to approximately 150 nmol ml^-1^. The bubble was removed and inhibitors were injected at this moment (10 μM DCMU, 10 μM MV, 10 μM DBMIB, 10 μM DNP-INT). Samples were illuminated via halogen lamp (Dolan Jenner, USA) at low light (120 μmol photons m^-2^ s^-1^) or high light (900 μmol photons m^-2^ s^-1^). At the end of each run the Chl concentration of each sample was determined in triplicate using the Porra Method (Porra, Thompson et al. 1989) in 90% MeOH to ensure accurate normalization of rates between samples. The cuvette was washed thoroughly with multiple rinses of 70% ethanol followed by MQ H_2_O when changing between inhibitors to avoid cross contamination. All data was analysed and fluxes calculated with equations described in (Beckmann, Messinger et al. 2009), which includes offsets for the changing relative concentrations of ^16^O_2_ and ^18^O_2_, whilst any background O_2_ consumption produced by thylakoid samples was normalised to zero in darkness as per (Furbank, Badger 1983).

### Gas exchange measurements in leaf discs

Gas fluxes from intact leaf discs in an atmosphere enriched in two stable isotopes, ^18^O_2_ and ^13^CO_2_, was performed using MIMS in an in-house built 1000 μl gas-phase cuvette connected to the same mass spectrometer, collecting masses 32, 36, 44 and 45 with a cycle time of approximately 6 seconds. 14 mm diameter leaf discs cut from fully developed WT *Arabidopsis thaliana* plants floated in darkness for 4 hours in a dish of H_2_O (control), H_2_O + MV (10 μM), H_2_O + DBMIB (20 μM) and H_2_O + DCMU (10 μM) at approximately 20 °C. For leaf disc samples infiltrated with catalase, SHAM and rotenone, discs were vacuum infiltrated before being allowed to dry on damp tissue in darkness for about one hour (until discs were no longer transparent). For all samples, in minimal light a 12.5 mm disc was cut from the incubated 14 mm discs, and loaded into the pre-calibrated cuvette at 25 °C. In darkness, the samples were purged with atmospheric air scrubbed of endogenous ^12^CO_2_ (Soda Lime, Li-Core, USA), the cuvette was then enriched to approximately 2% ^13^CO_2_ and 3% ^18^O_2_ by volume, with the remainder of the atmosphere comprising approximately standard atmospheric concentrations. The very low ^12^CO_2_ partial pressure in the beginning of measurements minimized membrane consumption of this isotope to almost zero, enabling the rate of its production in darkness (mitochondrial respiration) to be used as a basis for setting the background consumption rate of ^18^O_2_ (mitochondrial respiration in dark, plus Mehler reaction during illumination). Rates of ^16^O_2_ (PSII water splitting) and ^13^CO_2_ (CO_2_ fixation by Rubisco) were both set to zero in darkness. The enriched CO_2_ atmosphere avoided photorespiration, whilst an approximately atmospheric level of O_2_ was maintained to maximize the probability of observing Mehler associated O_2_ reduction, with the same offsets for steady dilution of the stable isotopes used in calculations of steady state fluxes as described for liquid phase measurements. Following initiation of data acquisition, samples experienced three minutes of darkness, five minutes of growth light (120 μmol photons m^-2^ s^-1^), five minutes of high light (900 μmol(photons) m^-2^ s^-1^) and then four minutes of darkness. Rates were calculated according to equations described in (Beckmann, Messinger et al. 2009).

## Acknowledgements

Research was funded by the Center of Excellence program of the Academy of Finland (project no 307335) and the Jane and Aatos Erkko Foundation.

## Refrences

Allen, J., 1977. Oxygen - a physiological electron acceptor in photosynthesis? Current Advances in Plant Science, 9, pp. 459–469.

Asada, K., 2006. Production and scavenging of reactive oxygen species in chloroplasts and their functions. Plant Physiology, 141(2), pp. 391–396.

Asada, K., 1999. THE WATER-WATER CYCLE IN CHLOROPLASTS: Scavenging of Active Oxygens and Dissipation of Excess Photons. Annual Review of Plant Physiology and Plant Molecular Biology, 50, pp. 601–639.

Baniulis, D., Hasan, S.S., Stofleth, J.T. and Cramer, W.A., 2013. Mechanism of enhanced superoxide production in the cytochrome b(6)f complex of oxygenic photosynthesis. Biochemistry, 52(50), pp. 8975–8983.

Bauer, R. and Wijnands, M.J.G., 1974. The Inhibition of Photosynthetic Electron Transport by DBMIB and its Restoration by p-Phenylenediamines; Studied by Means of Prompt and Delayed Chlorophyll Fluorescence of Green Algae. Zeitschrift fur Naturforschung - Section C Journal of Biosciences, 29(11-12), pp. 725–732.

Bechtold, U., Richard, O., Zamboni, A., Gapper, C., Geisler, M., Pogson, B., Karpinski, S. and Mullineaux, P.M., 2008. Impact of chloroplastic-and extracellular-sourced ROS on high light-responsive gene expression in Arabidopsis. Journal of experimental botany, 59(2), pp. 121–133.

Beckmann, K., Messinger, J., Badger, M.R., Wydrzynski, T. and Hillier, W., 2009. On-line mass spectrometry: membrane inlet sampling. Photosynthesis Research, 102(2-3), pp. 511–522.

Borisova-Mubarakshina, M., Ivanov, B.N., Vetoshkina, D.V., Lubimov, V.Y., Fedorchuk, T.P., Naydov, I.A., Kozuleva, M.A., Rudenko, N.N., Dall’osto, L., Cazzaniga, S. and Bassi, R., 2015. Long-term acclimatory response to excess excitation energy: evidence for a role of hydrogen peroxide in the regulation of photosystem II antenna size. Journal of experimental botany, 66(22), pp. 7151–7164.

Borisova-Mubarakshina, M.M., Naydov, I.A. and Ivanov, B.N., 2018. Oxidation of the plastoquinone pool in chloroplast thylakoid membranes by superoxide anion radicals. FEBS letters, 592(19), pp. 3221–3228.

Busch, F.A., Sage, T.L., Cousins, A.B. and Sage, R.F., 2013. C3 plants enhance rates of photosynthesis by reassimilating photorespired and respired CO2. Plant, Cell & Environment, 36(1), pp. 200–212.

Carpentier, R., 2001. The negative action of toxic divalent cations on the photosynthetic apparatus. In: M. Passarakli, ed, Handbook of plant and crop stress. New York: Marcel Dekker., pp. 763–772.

Cathcart, R., Schwiers, E. and Ames, B.N., 1983. Detection of picomole levels of hydroperoxides using a fluorescent dichlorofluorescein assay.

Chalier, F. and Tordo, P., 2002. 5-Diisopropoxyphosphoryl-5-methyl-1-pyrroline N-oxide, DIPPMPO, a crystalline analog of the nitrone DEPMPO: synthesis and spin trapping properties. Journal of the Chemical Society, Perkin Transactions 2, (12), pp. 2110–2117.

Cournac, L., Josse, E.M., Joët, T., Rumeau, D., Redding, K., Kuntz, M. and Peltier, G., 2000. Flexibility in photosynthetic electron transport: a newly identified chloroplast oxidase involved in chlororespiration. Philosophical transactions of the Royal Society of London.Series B, Biological sciences, 355(1402), pp. 1447–1454.

Crisp, P.A., Ganguly, D.R., Smith, A.B., Murray, K.D., Estavillo, G.M., Searle, I., Ford, E., Bogdanović, O., Lister, R., Borevitz, J.O., Eichten, S.R. and Pogson, B.J., 2017. Rapid Recovery Gene Downregulation during Excess-Light Stress and Recovery in Arabidopsis. The Plant Cell, 29(8), pp. 1836–1863.

Cui, F., Brosché, M., Shapiguzov, A., He, X., Vainonen, J.P., Leppälä, J., Trotta, A., Kangasjärvi, S., Salojärvi, J., Kangasjärvi, J. and Overmyer, K., 2019. Interaction of methyl viologen-induced chloroplast and mitochondrial signalling in Arabidopsis.

Dietz, K., Mittler, R. and Noctor, G., 2016. Recent Progress in Understanding the Role of Reactive Oxygen Species in Plant Cell Signaling. Plant Physiology, 171(3), pp. 1535–1539.

Durrant, J.R., Giorgi, L.B., Barber, J., Klug, D.R. and Porter, G., 1990. Characterisation of triplet states in isolated Photosystem II reaction centres: Oxygen quenching as a mechanism for photodamage.

Exposito-Rodriguez, M., Laissue, P.P., Yvon-Durocher, G., Smirnoff, N. and Mullineaux, P.M., 2017. Photosynthesis-dependent H(2)O(2) transfer from chloroplasts to nuclei provides a high-light signalling mechanism. Nature communications, 8(1), pp. 49–49.

Farquhar, G.D. and Busch, F.A., 2017. Changes in the chloroplastic CO2 concentration explain much of the observed Kok effect: a model. New Phytologist, 214(2), pp. 570–584.

Fichman, Y., Miller, G. and Mittler, R., 2019. Whole-Plant Live Imaging of Reactive Oxygen Species. Molecular Plant, 12(9), pp. 1203–1210.

Fitzpatrick, D., Aro, E. and Tiwari A. A., 2020. A Commonly Used Photosynthetic Inhibitor Fails to Block Electron Flow to Photosystem I in Intact Systems. Frontiers in Plant Science, 11, pp. 382.

Furbank, R.T. and Badger, M.R., 1983. Oxygen exchange associated with electron transport and photophosphorylation in spinach thylakoids.

Gollan, P.J. and Aro, E.M., 2020. Photosynthetic signalling during high light stress and recovery: targets and dynamics. Philosophical transactions of the Royal Society of London.Series B, Biological sciences, 375(1801), pp. 20190406.

Halliwell, B. and Gutteridge, J.M., 1984. Oxygen toxicity, oxygen radicals, transition metals and disease. The Biochemical journal, 219(1), pp. 1–14.

Heyno, E., Gross, C.M., Laureau, C., Culcasi, M., Pietri, S. and Krieger-Liszkay, A., 2009. Plastid alternative oxidase (PTOX) promotes oxidative stress when overexpressed in tobacco. The Journal of biological chemistry, 284(45), pp. 31174–31180.

Ilík, P., Pavlovič, A., Kouřil, R., Alboresi, A., Morosinotto, T., Allahverdiyeva, Y., Aro, E.M., Yamamoto, H. and Shikanai, T., 2017. Alternative electron transport mediated by flavodiiron proteins is operational in organisms from cyanobacteria up to gymnosperms. The New phytologist, 214(3), pp. 967–972.

Iwai, S., Ogata, S., Yamada, N., Onjo, M., Sonoike, K. and Shimazaki, K., 2019. Guard cell photosynthesis is crucial in abscisic acid-induced stomatal closure. Plant Direct, 3(5), pp. e00137.

Kaye, Y., Huang, W., Clowez, S., Saroussi, S., Idoine, A., Sanz-Luque, E. and Grossman, A.R., 2019. The mitochondrial alternative oxidase from Chlamydomonas reinhardtii enables survival in high light. The Journal of biological chemistry, 294(4), pp. 1380–1395.

Khorobrykh, A., 2019. Hydrogen Peroxide and Superoxide Anion Radical Photoproduction in PSII Preparations at Various Modifications of the Water-Oxidizing Complex. Plants (Basel, Switzerland), 8(9), pp. 329.

Khorobrykh, S.A. and Ivanov, B.N., 2002. Oxygen reduction in a plastoquinone pool of isolated pea thylakoids. Photosynthesis Research, 71(3), pp. 209–219.

Khorobrykh, S.A., Khorobrykh, A.A., Yanykin, D.V., Ivanov, B.N., Klimov, V.V. and Mano, J., 2011. Photoproduction of catalase-insensitive peroxides on the donor side of manganese-depleted photosystem II: evidence with a specific fluorescent probe. Biochemistry, 50(49), pp. 10658–10665.

Khorobrykh, S.A., Karonen, M. and Tyystjärvi, E., 2015. Experimental evidence suggesting that H2O2 is produced within the thylakoid membrane in a reaction between plastoquinol and singlet oxygen.

Khorobrykh, S. and Tyystjärvi, E., 2018. Plastoquinol generates and scavenges reactive oxygen species in organic solvent: Potential relevance for thylakoids.

Kimimura, M. and Katoh, S., 1972. Studies on electron transport associated with Photosystem I. I. Functional site of plastocyanin: Inhibitory effects of HgCl2 on electron transport and plastocyanin in chloroplasts.

Kitajima, S., Nii, H. and Kitamura, M., 2010. Recombinant stromal APX defective in the unique loop region showed improved tolerance to hydrogen peroxide. Bioscience, biotechnology, and biochemistry, 74(7), pp. 1501–1503.

Kozuleva, M.A., Ivanov, B.N., Vetoshkina, D.V. and Borisova-Mubarakshina, M., 2020. Minimizing an Electron Flow to Molecular Oxygen in Photosynthetic Electron Transfer Chain: An Evolutionary View. Frontiers in plant science, 11, pp. 211–211.

Lozier, R.H. and Butler, W.L., 1972. The effects of dibromothymoquinone on fluorescence and electron transport of spinach chloroplasts.

Malkin, R., 1981. Redox properties of the DBMIB—Rieske iron—sulfur complex in spinach chloroplast membranes.

Mehler, A.H., 1951. Studies on reactions of illuminated chloroplasts. I. Mechanism of the reduction of oxygen and other Hill reagents. Archives of Biochemistry and Biophysics, 33(1), pp. 65–77.

Mhamdi, A., Queval, G., Chaouch, S., Vanderauwera, S., Van Breusegem, F. and Noctor, G., 2010. Catalase function in plants: a focus on Arabidopsis mutants as stress-mimic models. Journal of experimental botany, 61(15), pp. 4197–4220.

Mubarakshina Borisova, M.M., Kozuleva, M.A., Rudenko, N.N., Naydov, I.A., Klenina, I.B. and Ivanov, B.N., 2012. Photosynthetic electron flow to oxygen and diffusion of hydrogen peroxide through the chloroplast envelope via aquaporins. Biochimica et biophysica acta, 1817(8), pp. 1314–1321.

Mubarakshina, M.M. and Ivanov, B.N., 2010. The production and scavenging of reactive oxygen species in the plastoquinone pool of chloroplast thylakoid membranes. Physiologia Plantarum, 140(2), pp. 103–110.

Nawrocki, W.J., Tourasse, N.J., Taly, A., Rappaport, F. and Wollman, F., 2015. The Plastid Terminal Oxidase: Its Elusive Function Points to Multiple Contributions to Plastid Physiology. Annual Review of Plant Biology, 66(1), pp. 49–74.

Noguchi, K. and Yoshida, K., 2008. Interaction between photosynthesis and respiration in illuminated leaves.

Porra, R.J., Thompson, W.A. and Kriedemann, P.E., 1989. Determination of accurate extinction coefficients and simultaneous equations for assaying chlorophylls a and b extracted with four different solvents: verification of the concentration of chlorophyll standards by atomic absorption spectroscopy. Biochimica et Biophysica Acta (BBA) - Bioenergetics, 975(3), pp. 384–394.

Rossel, J.B., Wilson, P.B., Hussain, D., Woo, N.S., Gordon, M.J., Mewett, O.P., Howell, K.A., Whelan, J., Kazan, K. and Pogson, B.J., 2007. Systemic and Intracellular Responses to Photooxidative Stress in Arabidopsis. The Plant Cell, 19(12), pp. 4091–4110.

Scarpeci, T.E. and Valle, E.M., 2008. Rearrangement of carbon metabolism in Arabidopsis thaliana subjected to oxidative stress condition: an emergency survival strategy. Plant Growth Regulation, 54(2), pp. 133–142.

Selinski, J. and Scheibe, R., 2019. Malate valves: old shuttles with new perspectives. Plant Biology, 21, pp. 21–30.

Shapiguzov, A., Nikkanen, L., Fitzpatrick, D., Vainonen, J.P., Gossens, R., Alseekh, S., Aarabi, F., Tiwari, A., Blokhina, O., Panzarová, K., Benedikty, Z., Tyystjärvi, E., Fernie, A.R., Trtílek, M., Aro, E., Rintamäki, E. and Kangasjärvi, J., 2020. Dissecting the interaction of photosynthetic electron transfer with mitochondrial signalling and hypoxic response in the Arabidopsis rcd1 mutant. Philosophical Transactions of the Royal Society B: Biological Sciences, 375(1801), pp. 20190413.

Shimakawa, G. and Miyake, C., 2018. Oxidation of P700 Ensures Robust Photosynthesis. Frontiers in Plant Science, 9, pp. 1617.

Telfer, A., Oldham, T.C., Phillips, D. and Barber, J., 1999. Singlet oxygen formation detected by near-infrared emission from isolated photosystem II reaction centres: Direct correlation between P680 triplet decay and luminescence rise kinetics and its consequences for photoinhibition.

Tiwari, A., Mamedov, F., Grieco, M., Suorsa, M., Jajoo, A., Styring, S., Tikkanen, M. and Aro, E.M., 2016. Photodamage of iron-sulphur clusters in photosystem I induces non-photochemical energy dissipation. Nature plants, 2, pp. 16035.

Tiwari, A. and Pospisil, P., 2009. Superoxide oxidase and reductase activity of cytochrome b559 in photosystem II. Biochimica et biophysica acta, 1787(8), pp. 985–994.

Villamena, F.A., 2017. Chapter 5 - EPR Spin Trapping. In: F.A. Villamena, ed, Reactive Species Detection in Biology. Boston: Elsevier, pp. 163–202.

Villamena, F.A. and Zweier, J.L., 2004. Detection of Reactive Oxygen and Nitrogen Species by EPR Spin Trapping. Antioxidants & Redox Signaling, 6(3), pp. 619–629.

Wang, W., He, E., Chen, J., Guo, Y., Chen, J., Liu, X. and Zheng, H., 2016. The reduced state of the plastoquinone pool is required for chloroplast-mediated stomatal closure in response to calcium stimulation. The Plant Journal, 86(2), pp. 132–144.

Witt, H.T., Rumberg, B., Junge, W. and Joliot, P., 1968. Electron Transfer, Field Changes, Proton Translocation and Phosphorylation in Photosynthesis, B. Hess and H. Staudinger, eds. In: Biochemie des, Sauerstoffs 1968, Springer Berlin Heidelberg, pp. 262–317.

Zhao, Y., Yu, H., Zhou, J., Smith, S.M. and Li, J., 2020. Malate Circulation: Linking Chloroplast Metabolism to Mitochondrial ROS. Trends in plant science, 25(5), pp. 446–454.

